# Deep networks may capture biological behaviour for shallow, but not deep, empirical characterizations

**DOI:** 10.1101/2022.03.02.482629

**Authors:** Peter Neri

## Abstract

We assess whether deep convolutional networks (DCN) can account for a most fundamental property of human vision: detection/discrimination of elementary image elements (bars) at different contrast levels. The human visual process can be characterized to varying degrees of ‘depth’, ranging from percentage of correct detections to detailed tuning and operating characteristics of the underlying perceptual mechanism. We challenge deep networks with the same stimuli/tasks used with human observers and apply equivalent characterization of the stimulus-response coupling. In general, we find that popular DCN architectures do not account for signature properties of the human process. For shallow depth of characterization, some variants of network-architecture/training-protocol produce human-like trends; however, richer empirical descriptors expose glaring discrepancies. These results urge caution in assessing whether neural networks do or do not capture human behaviour: ultimately, our ability to assess ‘success’ in this area can only be as good as afforded by the depth of behavioural characterization against which the network is evaluated. We propose a novel set of metrics/protocols that impose stringent constraints on the evaluation of DCN behaviour as adequate approximation of biological processes.

## 1. Introduction

Soon after the realization that DCN’s are capable of excellent performance in certain complex tasks, some authors have questioned whether the manner in which these networks perform such tasks mirrors the strategies adopted by humans or other biological systems [1, 2]. In other words, it is one thing to say that a given network is able to perform image classification efficiently; it is another thing to say that the behaviour exhibited by the network in achieving such high performance is an adequate approximation to the behaviour of a human observer performing the same task [3]. After all, birds fly and so do airplanes; but the wings of birds display movements that have little in common with those produced by aircraft devices.

In pursuing the above-stated question, current efforts in the literature have largely focused on expanding and/or complexifying what goes *into* the system [4]: the input stimuli with associated tasks (blue elements in **Figure 1**). At the same time, little effort has been devoted to a more thorough characterization of what comes *out* of the system: its behaviour. For example, early applications of machine learning tasks involved digit classification (input); the behaviour of the network (output) was evaluated using as metric the percentage of correct identifications [5]. In later applications, digit classification was extended to object recognition, generally regarded as a more complex task [6]; network behaviour, however, was still evaluated using essentially the same metric. There have been attempts at comparing machine with biological behaviour on a finer-grained scale [7, 8, 9], but they have been relatively uncommon and, for the purposes of the present study, fall under the same category as mainstream measures of overall performance.

**Figure 1.**
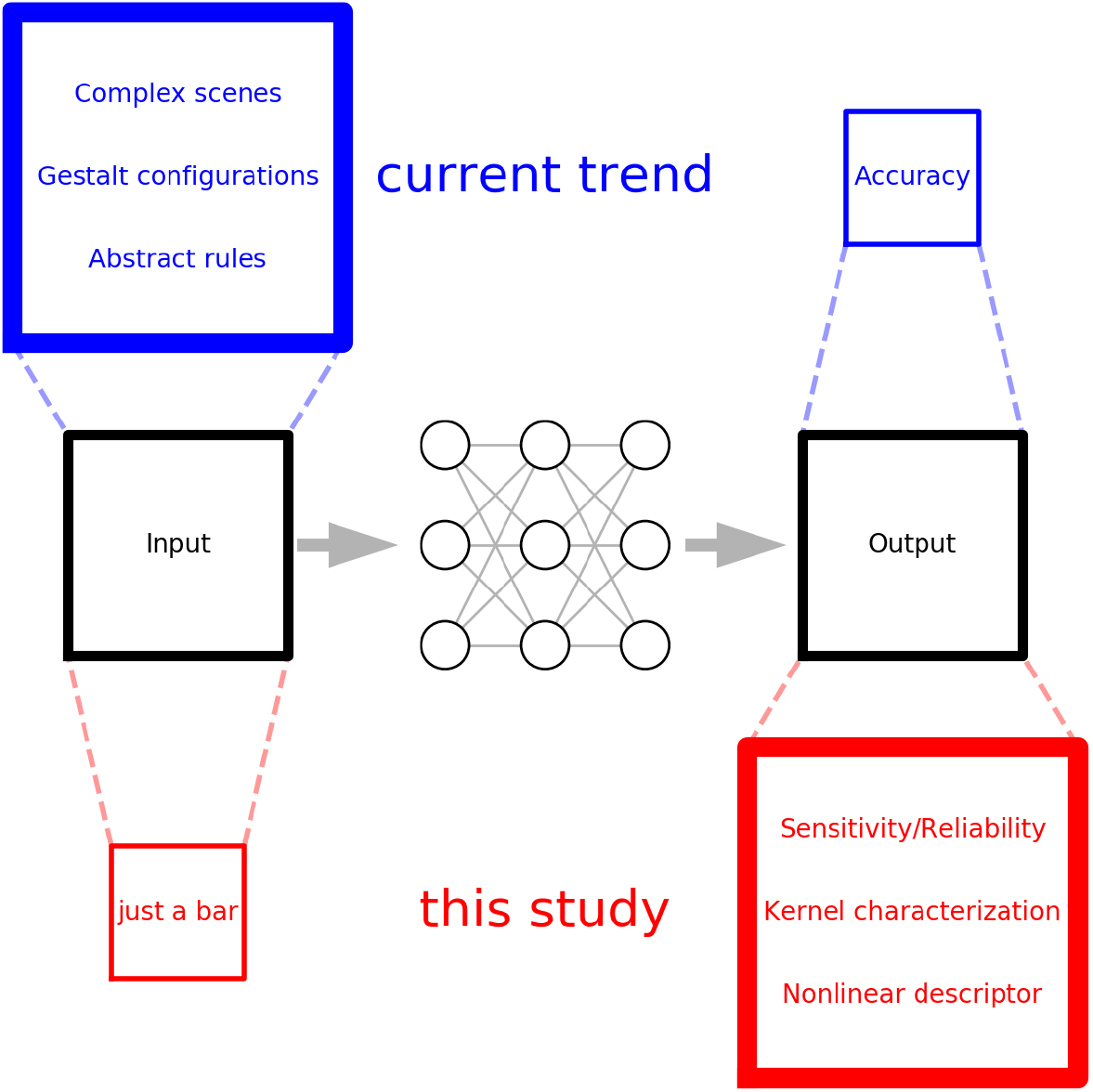
Looking at DCN’s from the other end. Current efforts towards exploring the applicability of DCN’s to human perception (blue elements) largely focus on expanding stimulus manipulations and task specifications (input) to discover potential discrepancies between humans and networks; examples include classification of complex scenes [4], counter-intuitive segmentation/grouping effects in contextual display [10], abstract relationships between objects and/or global spatial transformations [11]. Characterization of the output is mostly restricted to a simple performance measure such as area-under-the-curve (AUC). In this study, we take the opposite approach (red elements): the input is reduced to minimal specifications with associated tasks that are simplified to the essential operations of visual processing [12]; the output, on the other hand, is characterized to pain-staking degree via state-of-the-art behavioural tools that support a multi-faceted view of human/network functional architecture, comprising often overlooked properties such as intrinstic noise [13], linear transduction characteristics [14] and the potential contribution of dynamic nonlinearities [15].

In this article, we take the opposite approach (red elements in **Figure 1**). We deliberately simplify the input stimulus to a minimal design, while retaining a meaningful connection with the perceptual phenomenon of interest, namely human vision: instead of a complex natural scene, we present a vertical bar; in place of requiring that different breeds of dogs be correctly classified, we ask whether the bar is present or absent. At the same time, we strive to obtain a ‘deep’ characterization of the behaviour produced by both models and humans by capitalizing on recent developments in the quantification of animal perceptual processes [14, 15, 16]. The end result is a different way of looking at the potential relationship between neural networks and biological systems: there is no uncertainty associated with whether the input is represented adequately, because it is so simple that it could not be represented inadequately; the focus is shifted onto the idiosyncratic behaviour produced by the perceptual representation and its characterization, which is exploded into a far richer description. In a sense, we are looking at the problem from the other end (**Figure 1**).

When viewed from this inverted perspective, the problem at hand produces a number of interesting considerations. In particular, we find that substantially different conclusions may be reached regarding the biological plausibility of network models, depending on the degree of behavioural characterization against which machine-human alignment is evaluated. When the degree of characterization is comparable to that commonly adopted in contemporary applications, some network models may appear as adequate approximations to biological processes. However, when the output characterization is projected deeper into more articulate descriptors of the behavioural process, glaring discrepancies become evident between neural networks and the biological system they are meant to mimic, to the extent that the modelled behaviour may present opposite trends to those associated with the biological substrate. Our goal in demonstrating this outcome is not necessarily to prompt solutions for ameliorating the specific networks we used in relation to the specific stimuli/tasks they were challenged with, but rather to highlight more general issues associated with the complex task of determining whether artificial networks may serve as adequate descriptions of the behaviour produced by their biological counterparts [1, 2, 3].

## 2. Methods

### 2.1. Input/task specification

Stimuli consist of square images associated with binary labels, one label per image. Each image contains two arrays of 13 vertical bars: the ‘non-target’ array **s**^[0]^ and the ‘target’ array **s**^[1]^, placed to either side of the midline and separated by a small gap (**Figure 2C**). Each array is constructed by summing two components: a ‘noise’ component denoted by vector **n** and a ‘signal’ component denoted by vector **t**, so that **s**^[*q*]^ = **n**^[*q*]^ + **t**^[*q*]^ where *q* may denote either target (*q* = 1) or non-target (*q* = 0). Both **n** and **t** differ for target and non-target stimuli (hence the superscript *q*), however the difference is due to statistical sampling in the case of **n**, while it is deterministic in the case of **t**: **t**^[0]^ and **t**^[1]^ are pre-specified differently at the start of each experiment and remain unchanged throughout the experiment, while **n**^[0]^ and **n**^[1]^ are different samples from the same noise generator and differ not only between themselves, but also from one trial to the next.

**Figure 2.**
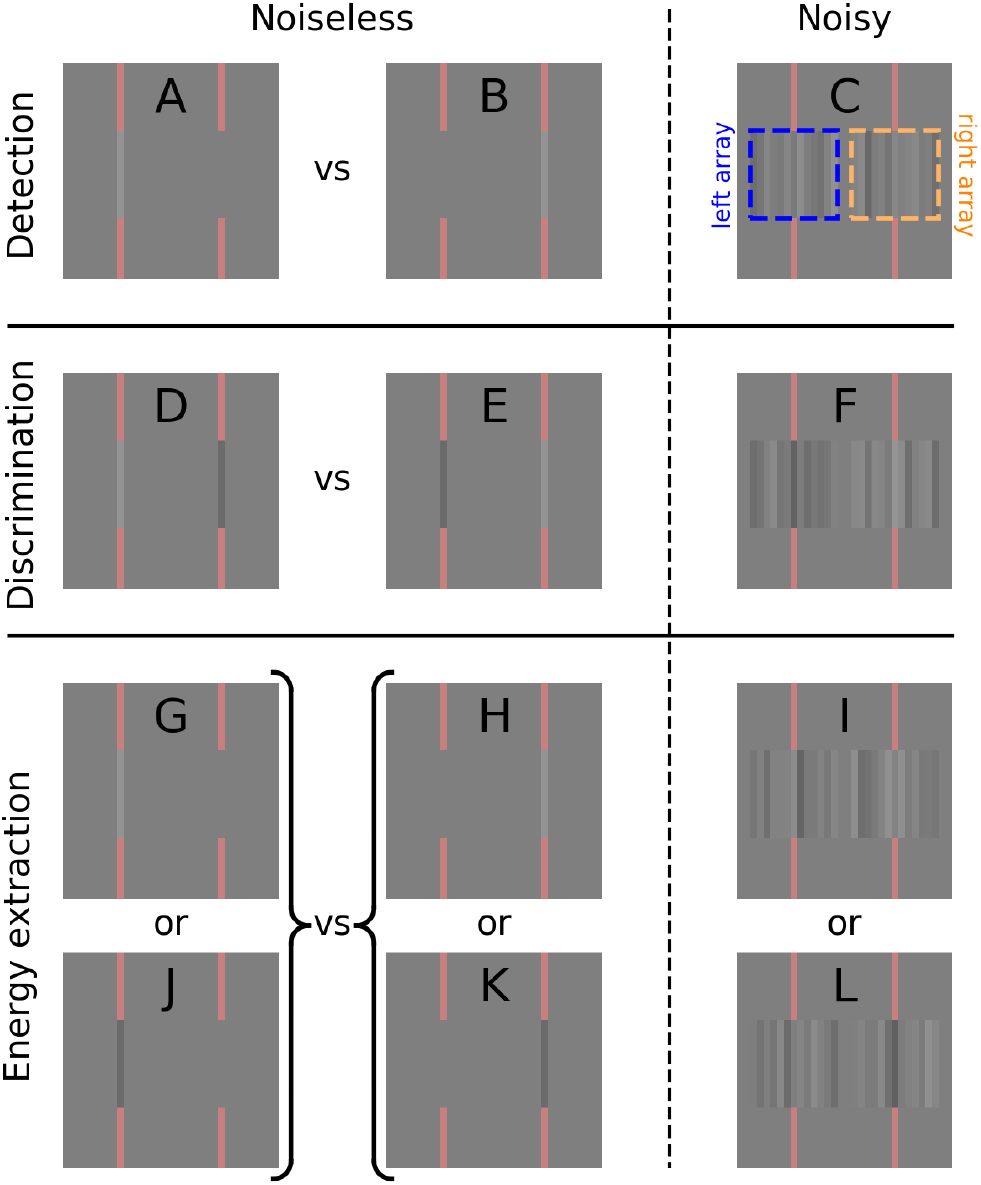
Stimuli and tasks. Each stimulus image (e.g. **C**) contains two one-dimensional arrays of 13 vertical bars each, one on the left and one on the right (the two arrays are indicated by blue/orange dashed rectangular outlines in **C**). In the detection task (**A**-**C**), one array contains a bright bar at its centre, while the other array is blank; the network is asked to determine whether the bar appeared within the array on the left (**A**) or within the array on the right (**B**). The two spatial positions potentially occupied by the target bar are indicated by red markers above and below the two arrays. Stimuli are rendered noisy by adding random intensity fluctuations to the different bars constituting each array (**C** shows an example obtained by adding noise to **B**). In the polarity-discrimination task (**D**-**F**), one array contains a bright bar while the other array contains a dark bar; the network is asked to determine which side contains the bright bar (**D** versus **E**). In the energy extraction task (**G**-**L**), one array contains a bar (randomly either bright or dark), while the other array is blank; the network is asked to determine whether the bar is on the left (**G**,**J**) or on the right (**H**,**K**). Notice that there is no labelling conflict across tasks: target images that appear in more than one task are associated with the same label.

Each element of **n** is independently and randomly drawn from a zero-mean Gaussian distribution with SD σ_N_, and **t** is zero everywhere except for the central element (corresponding to the seventh bar in the array) which takes value *τ*^[*q*]^: target signal **t**^[1]^ and non-target signal **t**^[0]^ only differ as specified by the two scalar quantities *τ*^[1]^ and *τ*^[0]^. The addition of target/non-target stimulus **t** therefore involves a simple luminance offset (*τ*) applied to the central bar of noise stimulus **n**. The different tasks involved detecting/discriminating offsets of this kind. Negative offsets involve a luminance decrement rather than an increment with respect to the background intensity of the image, which was fixed at 100 in the numerical simulations (corresponding background luminance was 30 cd/m^2^ in the human experiments). The spatial location of the central bar is marked on both sides by red vertical bars immediately above and below the arrays; these markers serve to minimize spatial uncertainty [17].

We consider three different tasks: detection, polarity discrimination, energy extraction. In the bright-bar ‘detection’ task, the target array contains a central bright bar of intensity *τ*^[1]^ > 0 added onto the noise, while the non-target array is just noise (*τ*^[0]^ = 0). If the target array appears on the left and the non-target array on the right (as in **Figure 2A**), stimulus label is ‘left’; if target array appears on the right and non-target array on the left (**Figure 2B-C**), label is ‘right’. In the ‘polarity discrimination’ experiments, *τ*^[1]^ > 0 and *τ*^[0]^ = −*τ*^[1]^: the target array contains a central bright bar while the non-target array contains a central dark bar of opposite intensity (**Figure 2D-F**). In the ‘energy extraction’ experiment, *τ*^[1]^ can be either positive (**Figure 2G-I**) or negative (**Figure 2J-L**) on any given trial by random assignment with equal probability, while *τ*^[0]^ = 0.

In the experiments with human observers [12], we also carried out a dark-bar variant (as opposed to bright-bar) of the detection task (*τ*^[1]^ < 0). Observers were specifically instructed to detect a dark bar (in the same way that they were specifically instructed to detect a bright bar for the bright-bar variant of the experiment), and the results were virtually identical to those obtained using a bright-bar target signal, except for the expected sign inversion of certain empirical descriptors (more specifically those of the first-order kind, to be detailed below). Because there is no conceptual difference between these two variants, we combine them into one detection dataset for the purposes of the present study; to merge the two datasets, we simply invert polarity of the dark-bar dataset and treat it as an extension of the bright-bar dataset.

We manipulate stimulus contrast and signal-to-noise ratio (SNR) as follows. *τ*^[1]^ can take one of 4 integer values between 1 and 4 when expressed as multiple of σ_N_; the assigned value is the stimulus signal-to-noise ratio (SNR). σ_N_ can be one of 4 logarithmically spaced values 1, 2, 4 or 8 in the numerical simulations (for the human experiments, this value is a multiple of physical monitor luminance of 0.3 cd/m^2^); the assigned value is the stimulus contrast level. For example, if SNR level is set to 2 and contrast level is set to 4 in the detection experiment, the stimulus is specified as follows: the non-target array consists of 13 bars, each taking a random value from a zero-mean Gaussian distribution with SD 4; the target array consists of 13 bars, each taking a random value from the same distribution, with a positive offset added onto the middle bar of 2 × 4=8. The two arrays are then added to the baseline image value of 100 (darker gray level in **Figure 2A**). Stimuli can therefore take one of 16 contrast/SNR combinations (4 contrast levels × 4 SNR levels).

### 2.2. Training/testing protocols

For the majority of the simulations, we trained networks on noiseless stimuli with high-intensity targets (e.g. **Figure 2A-B**): the intensity of the target bar was set to the maximum value afforded by the specifications above (32) and σ_N_=0. This was the protocol used with human observers when they were familiarized with the assigned stimulus/task before collection with noisy stimuli (such as the example in **Figure 2C**). All networks were then tested using independent noisy samples from the 16 configurations specified above. For a subset of the stimulations, we also trained on these 16 configurations, with and without added noise (those instances are explicitly indicated in the main text when discussing relevant results), however we obtained far poorer results using this approach (Appendix C). For this reason, noiseless high-intensity targets were the default option for training. This choice is also motivated by theoretical considerations (see Discussion).

Each stimulus mini-batch (size 64) was generated as follows. For detection/polarity-discrimination tasks, we generated 32 stimuli spanning all contrast/SNR configurations (16) and labels (2) with independent noise samples. We then generated another 32 stimuli of the same type, with inversion of target polarity for the energy-extraction task (no inversion was applied for the other two tasks). Stimuli were generated according to the specification associated with the given task (see above; see also **Figure 2**), and a given mini-batch could only contain stimuli/labels associated with one given task. We refer to a stimulus ‘ensemble’ as a sequence of 3000 mini-batches (therefore comprising 64×3000 stimuli). Ensembles can be either ‘blocked’, meaning that all mini-batches are associated with the same task, or ‘mixed’, meaning that different mini-batches can be associated with different tasks in random order. During training, a given stimulus ensemble was presented only once. During testing, a given stimulus ensemble was presented twice, so as to measure the proportion of instances on which the model produces the same response to the same stimulus on both occasions. We refer to this quantity as ‘double-pass agreement’. The figures specified above for mini-batch size and number of mini-batches were modified when required by a given training protocol (certain protocols either imposed specific sizes or prompted larger ones to allow for saturation of network learning trajectory).

Our network of reference is Lenet-5 (pyTorch implementation with final layer re-sized to binary output), for which stimulus dimension is 32×32×3; the information contained in each image of **Figure 2** can be rendered at this resolution without loss. We also test other architectures such as Alexnet (for which stimulus dimension is upsampled to 64×64×3) and Resnet-18 (224×224×3). In general, there is no appreciable improvement over Lenet-5. This is not surprising given the simplicity of our stimuli/tasks: despite being relatively small, Lenet-5 carries representational power well above that required to perform the assigned detection/discrimination tasks (consider that Lenet-5 is capable of adequately classifying CIFAR-10). We also verify that Lenet-5 is capable of producing a behavioural repertoire that spans the human range with relation to individual metrics. We explicitly address the more general issue of representational power in section 4.2.

Our training protocol is standard (stochastic gradient descent, cross entropy loss, learning rate of 0.001, momentum factor of 0.9, default pyTorch weight initialization). With the exception of transfer-learning applications (see below), all networks easily learn the assigned detection/discrimination tasks to optimal performance, so it is unlikely that training hyperparameters would impact our general conclusions. Because intrinsic noise is necessary to simulate behavioural variability as measured from humans (i.e. double-pass agreement<1), we implement it explicitly during the test phase. We introduce noise into the network by perturbing its weights using a fixed additive Gaussian source. The intensity of this source is selected so that it corresponds to a degree of output variability roughly compatible with that observed in humans. We verified that reasonable variations around these values have no impact on the results (by ‘reasonable’ we mean that they would not produce output variability well outside the range measured from human observers). When estimating human-model trial-by-trial agreement (metric introduced below), intrinsic noise is removed because we wish to maximize this quantity.

We trained randomly initialized networks on the detection/discrimination tasks, or we applied transfer-learning/finetuning to pre-trained configurations. In the latter procedure, we first train the network on the CIFAR-10/Imagenet databases and then reduce output size of the final layer to binary. When fine-tuning, we re-train all layers on the detection/discrimination tasks; when transfer-learning, we only re-train the final layer. With smaller networks (Lenet-5, CORnet-Z [18], scattering-transform with linear read-out [19], and a custom-made 3-layer network resembling a reduced version of Lenet-5), we carried out full training on the detection/discrimination tasks. With bigger networks (Alexnet, VGG-16 and Resnet-18 all pre-trained on ImageNet), we only applied transfer-learning/fine-tuning (we never trained the entire network from scratch). Our interest in transfer-learning the final layer stems from the notion that human observers, when asked to perform a novel visual task, may adjust flexible read-out modules involved in decision making, while relying on sensory signals from early circuitry that is relatively consolidated through lifetime experience [20, 21]. In other words, transfer learning may serve as a good approximation to human acquisition of the response rules involved in a novel sensory task (section 3.4).

### 2.3. Deep versus shallow characterization of sensory behaviour

We propose three different levels of characterization for the behavioural process, where ‘behaviour’ refers to the binary output of either human observer or neural network in connection with a specified input: a zeroth-order level of characterization based on classic performance metrics such as % of correct responses or d, sensitivity (green elements in **Figure 3**); a first-order level associated with the notion of linear filter, as in the perceptual template applied by human observers to the stimulus [22], or the overall feature selectivity of a convolutional layer (blue elements in **Figure 3**); a second-order level of characterization, essentially meant to capture behaviour that is not adequately represented by first-order descriptors (red elements in **Figure 3**). Before discussing specific issues of interpretability (or lack thereof) associated with these different elements, we attempt to develop an intuitive understanding for their general significance in relation to the problem at hand.

**Figure 3.**
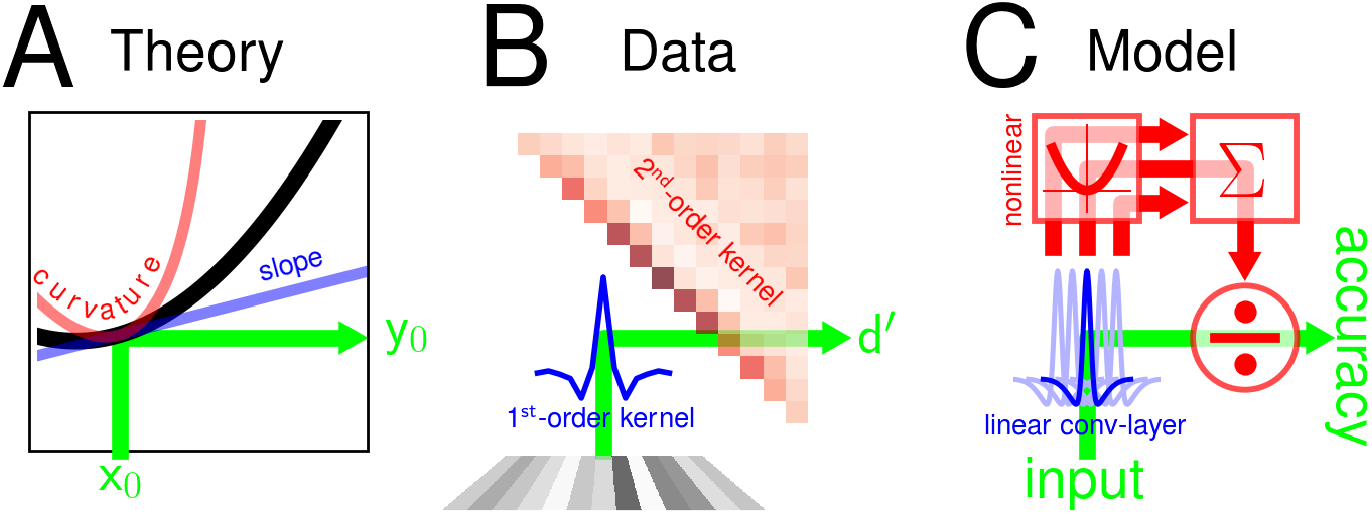
The behaviour of a human observer or a neural network can be characterized to different degrees of depth, similar to the way in which a simple function can be approximated to different orders of a Taylor expansion (**A**). For systems such as human observers, the corresponding characterization may be limited to sensitivity (d,) or may be extended to perceptual kernels of the first- and second-order [23] (**B**). Under some conditions, it is possible to establish a connection between first-order kernels and the linear portion of small network models such as convolutional layers (blue in **C**), as well as between second-order kernels and nonlinear model components such as full-wave rectifiers (red in **C**), however the link is not transparent [16].

There are at least three complementary ways of understanding the different levels outlined above. The most familiar approach is to draw a parallel with Taylor expansion (**Figure 3A**). This parallel is not entirely accurate, but it does retain a meaningful connection with tools that are routinely used in nonlinear system characterization [15]; more specifically, the output generated by a functional system such as a neuron, a human observer or a network model, can be described via Volterra expansion [24, 25] (**Figure 3B**). In a sense, Volterra kernels are generalized Taylor coefficients. For the specific case of the stimulus considered here, we may approximate the scalar response of the system to one stimulus array as a quadratic projection *r*^[*q*]^ = *h*_0_ + ⟨**s**^[*q*]^, **h**_1_⟩ + ⟨**s**^[*q*]^ ⊗ **s**^[*q*]^, **h**_2_⟩ (⟩, ⟩ is (Frobenius) inner product, ⊗ is outer product, **h**_2_ is a matrix); the system then produces label 0 if *r*^[0]^ > *r*^[1]^, label 1 otherwise (output response rule). Because the binary response rule renders *h*_0_ (constant baseline term) irrelevant, a zeroth-order characterization of this system is not directly connected with the zeroth-order coefficient in a Taylor expansion. We regard the parallel with Taylor expansion as a useful framework for intuitive purposes, with the understanding that it does not capture important aspects of the tools we introduce below.

The second viewpoint is central to this study: it involves tools for capitalizing on the output response to different degrees of characterization (**Figure 3B**). At the most basic level (zeroth order) we simply measure performance as proportion of correct responses *p*_c_ (transformed to Z units and therefore expressed as d′ for unbiased responses), a standard metric of behavioural sensitivity connected with AUC [26]. We also measure a related quantity, which we term ‘agreement’ (*p*_a_): the proportion of responses that are replicated by the system when an identical stimulus sequence is presented a second time [27, 28, 29]. A purely deterministic system (e.g. a pre-trained network with no internal noise at the time of testing) would produce an agreement of 1 (same response to both passes on every instance), even though the % of correct responses may be less than 100%. The presence of internal variability (i.e. intrinsic to the system as opposed to stimulus-driven) causes agreement to drop below 1, potentially with no concomitant change in *p*_c_. In this sense, *p*_c_ and *p*_a_ are related quantities but not entirely redundant: they do carry independent information for the purpose of system characterization [29].

Mainstream applications of signal detection theory, in both human psychophysics and machine learning, largely rely on the level of characterization detailed above (although the relevant metrics are often used parametrically to derive more complex descriptors, such as just-noticeable-increments and thresholds). Here we push output characterization further. At the next level, we compute first-order descriptors via reverse correlation of the input noise with the output response [30, 14, 15]. More specifically, we classify each input noise sample as **n**^[*q,z*]^, where *z* = 0 (1) if the system produces the correct (incorrect) label for the corresponding stimulus. For example, **n**^[1,1]^ is the noise sample that is added to the target array (*q* = 1) for a stimulus which the system labels correctly (*z* = 1). The first-order descriptor consists of two sub-components: the *target-present* descriptor 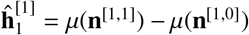 where µ is average across samples of the indexed type; and the *target-absent* descriptor 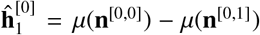. The full first-order descriptor is simply 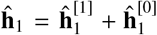. The first-order descriptor **ĥ**_1_ is therefore obtained by *averaging* noise samples [30].

We can achieve a further level of characterization by extending the above-detailed approach to the ‘second-order’ descriptor **ĥ**_2_, derived from the *covariance* of noise samples [23, 31, 15]: **ĥ**_1_ = (2δ_*q,z*_ − 1) ∑_*q,z*_ cov(**n**^[*q,z*]^) where δ_*q,z*_ is Kronecker δ (=0 if *q* ≠ *z*, =1 if *q* = *z*). This expression is equivalent to the one for **ĥ**_1_ except µ has been replaced by covariance matrix cov. In the notation above, we draw an important distinction between **h**_1_/**h**_2_ and **ĥ**_1_/**ĥ**_2_ [32]: the former is a theoretical construct (kernel), the latter is an experimental object (descriptor). The connection between the two is not trivial [16].

The third perspective is informed by network architecture: it is centred around computational circuits that may be connected with the empirical characterization detailed above (**Figure 3C**). The only circuit for which this connection is unproblematic is the linear-nonlinear (LN) cascade [14]. In the human literature, this model is often termed ‘template-matcher’ [22]: *r*^[*q*]^ = ⟨**s**^[*q*]^, **w**⟩ where **w** is the template that is linearly matched against the stimulus; the nonlinear operation is represented by the output response rule (see above). In the context of machine learning applications, the simplest implementation is the Perceptron [33]. For this class of models, it is expected that **ĥ**_1_ ∝ **w** and **ĥ**_2_ = 0 [15]. This prediction is neat, however its utility is limited by the applicability of the LN model: in both human psychophysics and machine learning, the LN cascade is useful as a normative model (e.g. ideal observer theory), not as a realistic/powerful application [34, 35]. Departures from the LN architecture will be reflected by structured modulations within **ĥ**_1_ and **ĥ**_2_, which can then be exploited for the purpose of constraining candidate circuits [15]; having said that, there is no *transparent* connection between specific circuit components and specific kernel modulations [16].

The power and appeal of the approach in **Figure 3B** derive from its applicability to both humans and neural networks under virtually identical conditions. For the class of minimal stimulus/task specifications adopted here, there is no room for ambiguity as to how the external noise source should be structured, nor as to how it can be effectively exploited to maximize characterization of the system. In this study, we capitalize on this point of contact between human psychophysics and machine learning to gain some insight into how this connection can be better understood, particularly with relation to potential pitfalls that may be associated with it.

## 3. Results

### 3.1. Network-vs-human comparison across three orders of characterization

Our core interest is behaviour as a dynamic process: we want to understand how behaviour is modified by specific stimulus manipulations. For this reason, we are primarily interested in the *trend* displayed by changing values of a given behavioural metric as a function of changing configurations for the input stimulus, rather than the *absolute* value of the metric for a specific stimulus configuration. In other words, it is far more informative to know that e.g. sensitivity increases with increasing contrast, than knowing that sensitivity is 1.2 for a contrast value of 9%: the latter property will surely depend on unimportant details of the stimulus, whereas the former property is likely to generalize across different stimulus configurations and possibly tasks, therefore exposing an important structural feature of the underlying mechanism. We therefore focus on how a given behavioural metric varies as a function of the 4×4 contrast×SNR stimulus configurations detailed above (section 2.1), without much regard for the absolute values of said metric (provided they remain within a reasonable/plausible range).

The different panels in **Figure 4B** plot human sensitivity (radius of green circle) as it varies with increasing contrast (left to right panels) and increasing SNR (bottom to top panels). As expected, there is marked improvement in sensitivity with SNR (top circles are larger than bottom circles); there is also milder improvement with contrast (right-hand circles are slightly larger than left-hand circles). **Figure 4F** plots corresponding data for Lenet-5. For this figure, the entire network was trained only on the detection task with noiseless stimuli; under these conditions, it easily learns the task (**Figure 4E**). It is clear that, at least qualitatively, the dependence of network sensitivity on contrast and SNR mirrors the human trend: absolute values are different (network sensitivity is overall higher) and the effect of contrast is more marked for the network, but the same general tendency is observed for both human and model (compare **Figure 4B** with **Figure 4F**).

**Figure 4.**
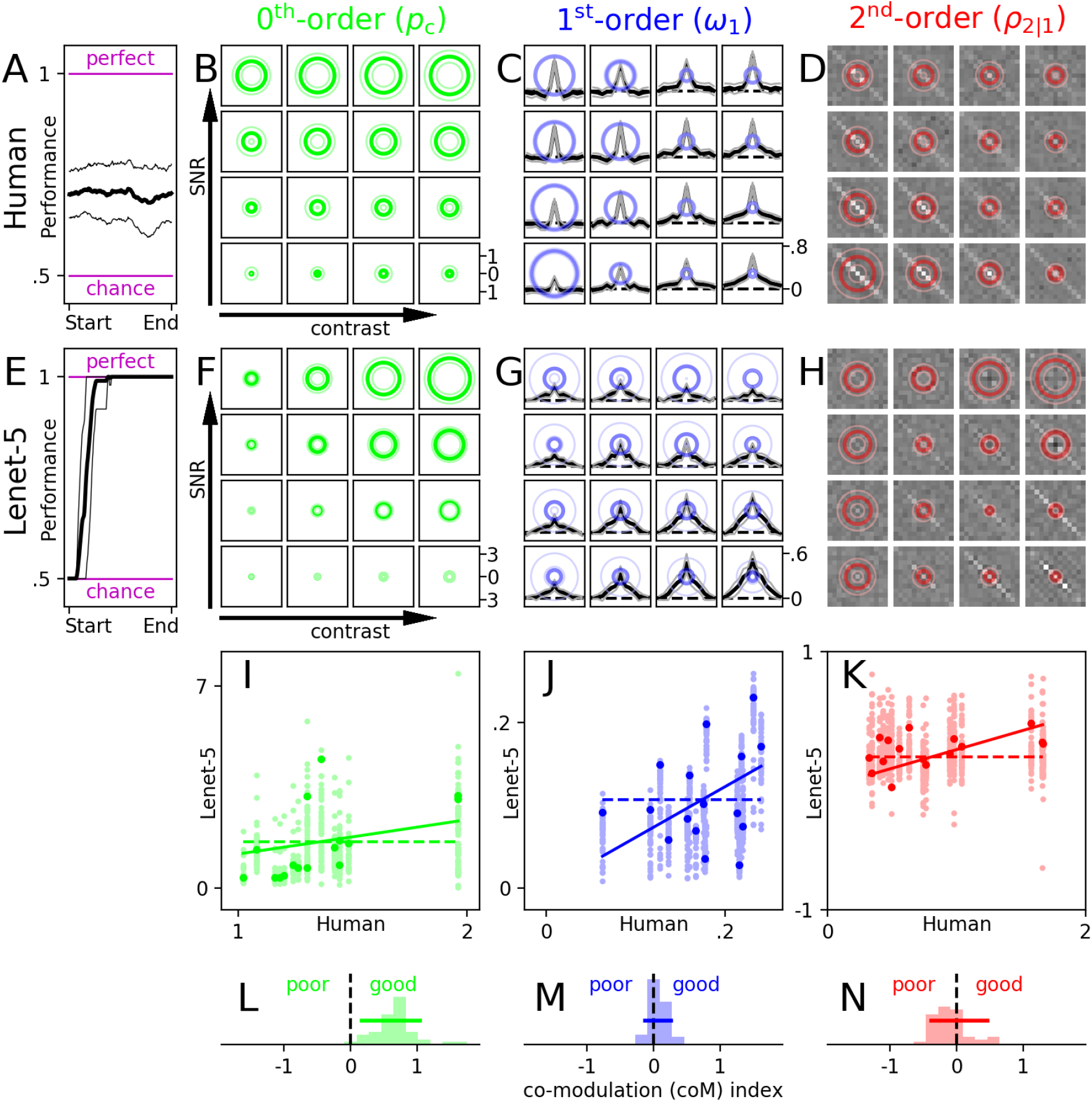
Lenet-5 captures zeroth-order human behaviour, but not higher orders. Performance (% correct) in detection task (**Figure 2A-C**) across consecutive blocks from each human participant is down-sampled to number of blocks collected by participant with least amount of data (241 blocks) to align start/end of data collection and allow averaging across participants. Resulting trace is smoothed with moving window (length=50) and plotted in **A** (thin lines show ±1 SD across participants). Humans demonstrate no measurable perceptual learning (performance remains stable across data collection); stimulus parameters were intentionally selected to target threshold-level performance (neither chance nor perfect) as required for the application of reverse correlation [30, 14]. **E** plots average performance across mini-batches during training of Lenet-5 with noiseless high-contrast/high-SNR stimuli (thin lines show ±1 SD across different simulations); this network consistently learns the detection task (100% correct at end of training). Radius of circles in **B** plots sensitivity (Z-transformed % correct) across 4 levels of stimulus contrast and 4 levels of stimulus SNR (hence the 4×4 grid); thick line shows mean, thin lines show ±20 SEM. Black traces in **C** plot first-order descriptors (thin lines show ±2 SEM); radius of blue circle plots sharpness (spectral centroid) for corresponding trace (thin lines show ±1 SEM). Surface plots in **D** show second-order descriptors (bright for positive, dark for negative); radius of red circle plots second-order/first-order RMS ratio for corresponding descriptors (thin lines show ±10 SEM). **F**-**H** plot same as **B**-**D** for Lenet-5; thin lines now show ±1 SD across simulations. **I** plots the 16 senstivity values from **B** on x axis versus the corresponding 16 values from **F** (y axis) for 50 different network simulations; the outcome of a representative simulation is highlighted by larger more saturated symbols, alongside associated best-fit scaling relationship (solid line) and constant relationship (dashed line). A co-modulation index is computed for each simulation by taking log-ratio between MSE’s associated with these two relationships (see main text); the distribution of this index across simulations is plotted in **L** with 5^th^-95^th^ percentile range indicated by horizontal segment. **J**,**M** plot same for sharpness, **K**,**N** for second-order/first-order RMS ratio.

As explained above, our focus is on trends, and more specifically on comparing trends between human and network. We quantify the similarity between trends via the ‘co-modulation index’, computed as follows: we plot human versus corresponding network sensitivity values for each of the 16 stimulus configurations (denoted by vectors **x**_*human*_ and **x**_*net*_, plotted on x and y axes in **Figure 4I**); we find the best-fitting scaling coefficient *k* such that the MSE between *k***x**_net_ and **x**_human_ is minimized (the corresponding linear fit is shown by the solid line in **Figure 4I**); we compute the RMS of the difference between **x**_human_ and *k***x**_net_ and call it ΔRMS; we compute the RMS of the difference between **x**_human_ and its average value (indicated by the dashed line in **Figure 4I**) and call that ΔRMS_0_; the co-modulation index is log(ΔRMS)−log(ΔRMS_0_). In words, the co-modulation index quantifies the extent to which simulated results show the same dependence on SNR/contrast as the human results with reference to a baseline hypothesis of no dependence, the latter corresponding to a fixed value (average estimate) across all 16 stimulus configurations. Besides effectively encapsulating the essence of the notion we wish to measure (see below), this metric can be computed from a simple analytical expression.

To motivate our choice of the above metric, we ask the following question: why not simply use the standard Pearson correlation coefficient? The co-modulation index detailed above only allows for rescaling (the same index is obtained for **x**_*net*_ and *g***x**_*net*_ where *g* is an arbitrary rescaling factor), whereas the correlation coefficient allows for both rescaling and shifting (the same coefficient is obtained for **x**_*net*_ and *g***x**_*net*_ + *s* where *s* is an arbitrary additive factor). The reason for not incorporating an additive shift into the co-registration between human and model data is that it can lead to an overall change of sign, and this corresponds to a qualitatively different meaning for the metrics we adopt here. In the case of Z-transformed proportion of correct responses (*p*_c_), for example, 0 carries a specific meaning: chance performance. It is therefore unwarranted to let simulated values cross the 0 point: this may lead to a better match with the human data (e.g. if ‘match’ is quantified via correlation coefficient), but it involves re-assigning a new (different) meaning to the simulated results, therefore distorting interpretation of the distance metric. On the contrary, re-scaling does not alter the meaning of the metric, and an overall change of scaling factor can be effected without modifying the architecture of the system: for example, a set of *p*_c_ values can be uniformly amplified by simply reducing internal noise [26]. The co-modulation index is designed to factor out parameters of this kind, while instead emphasizing invariant trends that are reflective of system architecture. Finally, we have verified via visual inspection of several simulations that the co-modulation index provides a meaningful reflection of the qualitative match between human and network descriptors, while correlation coefficients sometimes produce awkward results.

**Figure 4L** plots the distribution of the Human-vs-Lenet-5 co-modulation index for sensitivity as it varies across 50 simulations of the network. Clearly this index remains positive (green distribution falls to the right of the vertical dashed line indicating 0), despite fluctuations from simulation to simulation, consistent with our earlier qualitative comparison between **Figure 4B** and **Figure 4F**. In the remaining part of this study, we consider a result of this kind as an indication of ‘good’ match between human observers and network simulations, even though the range spanned by the co-modulation index may be smaller than it is in **Figure 4L**: as long as the index remains positive, we regard the neural network as capturing at least some of the architectural features associated with the human perceptual mechanism. Rather than detailing numeric assessment of individual effects (this would produce an unwieldy amount of textual details given the large number of architectures/protocols/conditions/metrics examined in this study), we adopt a visual indicator of data spread (percentile range indicated by horizontal line **Figure 4L**).

The black traces in **Figure 4C** and **Figure 4G** plot first-order kernel descriptors (**ĥ**_1_) as a function of contrast/SNR using the same 4×4 panel arrangement adopted for sensitivity in **Figure 4B,F**. Qualitative inspection of these traces suggests that the manner in which their shape varies as a function of contrast/SNR is different between humans and network: for human observers, traces are bandpass at low contrast (left-most panels in **Figure 4C**) and lowpass at high contrast (right-most panels); for the neural network, contrast only appears to scale the amplitude of the traces at low SNR (lower panels in **Figure 4G**), without substantially altering their shape. We would like to capture these observations in the same format we used for the zeroth-order characterization. This is essentially an exercise in data reduction (see immediately below).

The first step involves converting each trace into a meaningful scalar value, one per contrast/SNR stimulus configuration. A suitable map is the spectral centroid, a measure of trace ‘sharpness’ [12]: we compute the power spectrum **p** of trace **ĥ**_1_ across frequency values **f**; we normalize **p** to unit-sum, and rescale **f** to range between 0 (DC) and 1 (maximum frequency); the spectral centroid ω_1_ is ⟨**p, f**⟩. This quantity is indicated by the radius of the blue circles in **Figure 4C**: in line with the qualitative description above, sharpness is high (large circles) at low contrast (left-most panels), and low (small circles) at high contrast (corresponding to the contrast-dependent highpass-to-lowpass transition we documented in previous work [12]). This human pattern is not reflected by the network-generated pattern of blue circles in **Figure 4G**; consequently, the co-modulation index for this metric is centred around 0 (**Figure 4M**).

The surface plots in **Figure 4D** and **Figure 4H** show second-order kernel descriptors (**ĥ**_2_). As with first-order descriptors, qualitative comparison between human and network is sufficient to expose noticeable differences in their patterns of behaviour. In humans (**Figure 4D**), second-order descriptors present substantial diagonal modulations at low contrast (left-most panels) and little or no structure at high contrast (right-most panels); for Lenet-5 (**Figure 4H**), most second-order modulations are instead observed at high contrast and low SNR (lower-right panels).

In mapping each 2D-surface descriptor onto a scalar value, we consider that the amplitude of the second-order descriptor is best compared in relation to the amplitude of the first-order descriptor, rather than on its own as an absolute quantity: for a system with comparable contributions from linear and nonlinear processing components (**Figure 3C**), the absolute amplitude of a given descriptor (whether first- or second-order) is expected to depend on non-architectural factors such as internal noise, while the amplitude ratio between descriptors is expected to remain more stable in the face of changes associated with said factors [15]. In other words, amplitude ratio is a better index of system linear/nonlinear architecture [35]. We therefore compute the log-ratio between the RMS of the second-order descriptor and the RMS of the first-order descriptor ρ_2|1_ = log(|**ĥ**_2_|)−log(|**ĥ**_1_|) where |**x**| is RMS of vector/matrix **x**; this quantity is indicated by the radius of the red circles in **Figure 4D,H**. Again, we notice poor alingment between human and network across the 4×4 panels: if anything, there appear to be opposite trends. This impression is confirmed by quantification via the co-modulation index, which shows a tendency towards negative values (**Figure 4N**).

To summarize the above, we find that Lenet-5 seems like a promising candidate for modelling the human process based on a zeroth-order characterization of human/network behaviour. However, this assessment must be revised at the first order of characterization, and even reversed at the second order: the network goes from good to poor to bad. We now proceed to demonstrate the generality of this pattern by adopting different training protocols and network architectures.

### 3.2. The co-modulation plot

Before proceeding on to an extensive exploration of network parameterization and training diet, we want to develop a tool for quickly evaluating the viability of a specific network such that the tool successfully trades off ease of inspection with metric richness. We start by expanding each order of characterization with the addition of a complementary metric, so that the 3 metrics detailed above (sensitivity, first-order sharpness, second-to-first-order amplitude ratio) are augmented by 3 more related metrics; we then reduce the resulting high-dimensional characterization into one plot that retains all essential information and enables an informed judgment as to the viability of a given network.

The complementary metric we adopt for the zeroth-order characterization is (Z-transformed) *p*_a_, introduced above. This metric provides an indirect measure of system reliability in the sense of reproducibility: it reflects the amount of noise intrinsic to the system, i.e. the portion of output variability that is decoupled from the stimulus [28]. Together with *p*_c_, these two measurements are sufficient to constrain parameterization of a signal-detection-theory (SDT) model of the sensory process [29]. The complementary metric for the first-order characterization is simply RMS amplitude of the first-order descriptor |**ĥ**_1_|, in line with the commonly adopted approach of capturing variations in linear filter shape via separation into two components: tuning change (sharpness) and amplitude change (RMS) [36, 37]. For the second-order characterization, we obtain a complementary estimate of the extent to which the system is nonlinear as opposed to linear via 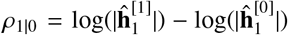; in words, this is the log-ratio between the RMS of the target-present first-order descriptor 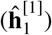 and the RMS of the target-absent descriptor 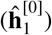. It is a well-established metric of departure from the template-matcher [38, 39, 35], and its theoretical connection with the second-order descriptor is well understood [23, 15].

**Figure 5A** plots the co-modulation index for a primary metric (e.g. *p*_c_) on the x axis versus the corresponding co-modulation index for the complementary metric (e.g. *p*_a_) on the y axis, across all network simulations (one data point per simulation) and for the three different levels of characterization (different colours). This device, which we term co-modulation plot, has been specifically designed to allow estimation of all relevant effects ‘at a glance’. It is important to this visualization tool that the quantity plotted on both axes is *not* the metric itself (e.g. *p*_c_), but its co-modulation index: a measure of the similarity between human and network with relation to *trends* of variation for a given metric as a function of input contrast/SNR manipulations (see previous section). Furthermore, the comodulation index is re-scaled separately for each metric (e.g. *p*_c_) to range between -1 and 1. This normalization is applied for two reasons: first, to make different metrics/orders more directly comparable; second, because this procedure effectively emphasizes the degree of co-modulation *spread* for a given metric with respect to the 0 point, regardless of absolute value. For example, if a given co-modulation index is generally small and positive, as long as it is *consistently* positive, it will be emphasized by this procedure.

**Figure 5.**
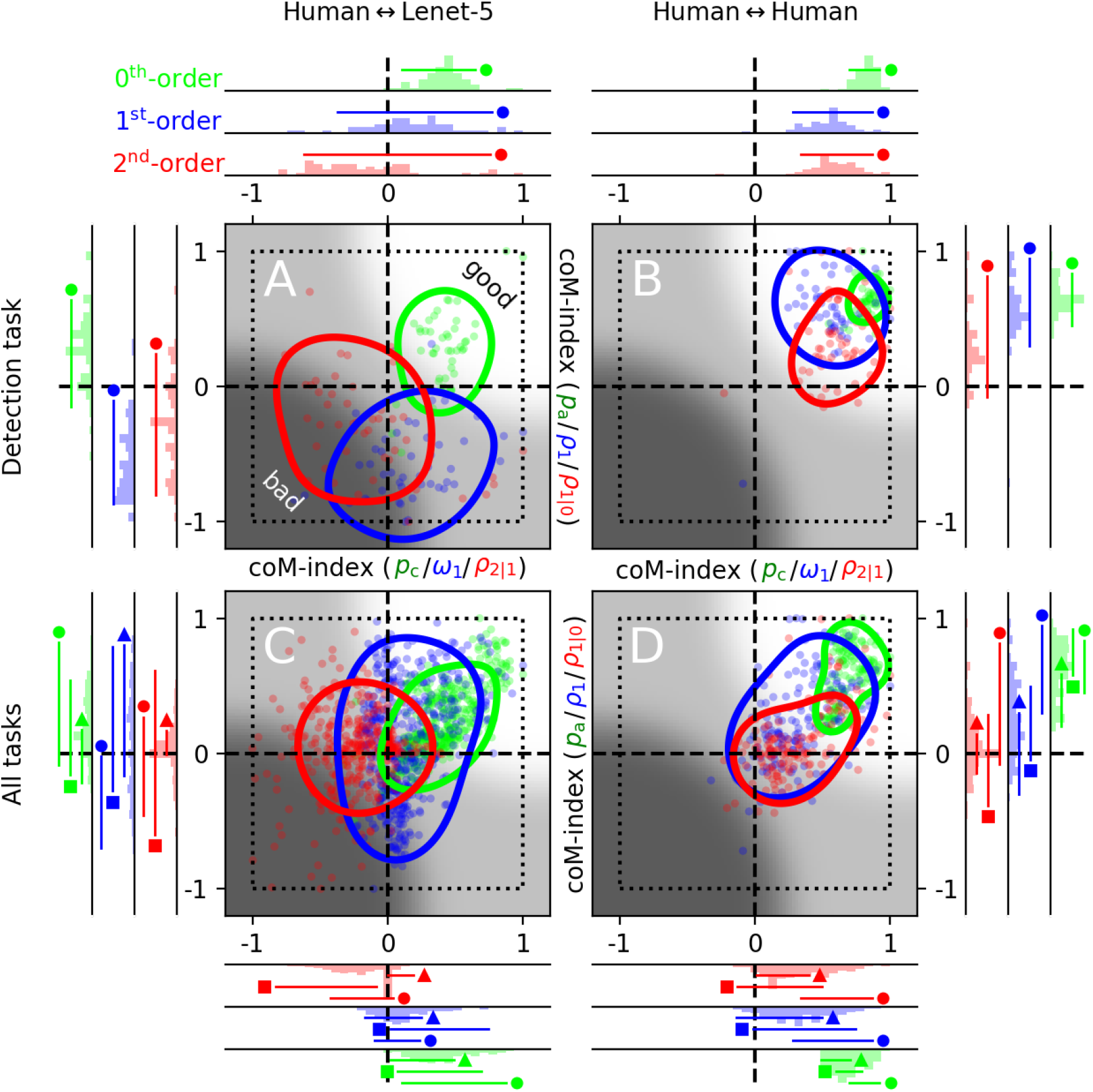
Co-modulation plots: visual tools for model evaluation ‘at a glance’. Abscissa values in **A** plot co-modulation indices from **Figure 4L-N**, separately rescaled for each metric to range between -1 and 1. Each of these primary metrics is associated with a complementary metric (whose co-modulation index is plotted on the y axis): % correct (x coordinates of green data points) is associated with % agreement (y coordinates of green data points); sharpness of first-order descriptor (x coordinates of blue data points) is associated with its RMS (y coordinates of blue data points); second-order/first-order RMS log-ratio (x coordinates of red data points) is associated with target-present/target-absent log-ratio of first-order RMS (y coordinates of red data points). **B** is plotted to the same conventions, but shows co-modulation indices between half the human dataset (odd trials) and the remaining half (even trials). **C**-**D** combine data from all three tasks; although the distinction between different tasks is not retained for the scatter plot, it is maintained at the level of the marginal histograms: •is placed next to the line marking 5^th^-95^th^ percentile range for detection, ■ for polarity-discrimination, ▴ for energy-extraction. For **C**, Lenet-5 is trained on all 3 tasks simultaneously. Each dot in **A**,**C** (network) refers to a different iteration of the same simulation; each dot in **B**,**D** (human) refers to a different bootstrap sample from the experimental dataset. Contours slice through blurred data distribution (data points convolved with Gaussian weighting function with SD equal to 0.4 of plot size) at 1/3 of peak height; they reflect data spread and are only intended for visualization purposes.

In the co-modulation plot, a good match between human and network corresponds to data falling within the upper-right quadrant (positive co-modulation index), away from the origin. This region of ‘viability’ is indicated by brighter background. Other regions of the co-modulation plot (dark background) correspond to network failures in terms of capturing human behaviour. It is clear from looking at this plot that the zeroth-order description falls within the ‘good’ region (green data points scatter within the bright area), while the other two orders of description fall within the ‘bad’ region (blue/red data points scatter within the dark area), consistent with our earlier evaluation based on **Figure 4**.

There is a potential pitfall with the approach detailed above, in that there are two possible reasons why the comodulation index may not be significantly different than zero. Under one scenario, the metric pattern produced by the network lacks structure or, if it carries structure, said structure is orthogonal to the human pattern. Under an alternative scenario, it is the metric pattern produced by human observers that lacks structure, i.e. it only reflects measurement noise. Under the latter scenario, human metric measurements are uninformative about the human process and are therefore equally uninformative about the potential viability of the network. To determine which one of these two scenarios is applicable for a given metric, we compute the corresponding human-vs-human co-modulation index in place of the human-vs-network co-modulation index: instead of pitting the entire human dataset against the network, we pit half the human dataset (odd trials) against the remaining half (even trials).

If the human measurements carry structure, we expect positive human-vs-human co-modulation indices. It should be noted that if the co-modulation index is *not* significantly different than 0, this does not necessarily imply that the human dataset does not carry any measurable structure: the human-vs-human co-modulation index is effectively computed from half the original dataset and is therefore expected to carry less resolving power than the entire human dataset. However, we allow this implication to carry through here in an effort to maintain a conservative approach towards model evaluation. It is clear from the human-vs-human co-modulation plot in **Figure 5B** that, for the detection task we have been examining so far, all metric measurements carry structure (all data points fall within bright region). This implies that human-vs-network co-modulation index values that scatter around 0, such as the sharpness index for the first-order characterization (x value of blue data points in **Figure 5A** and distribution in **Figure 4M**), indicate a genuine failure on the part of the neural network to capture structured patterns of human behaviour.

**Figure 5D** combines human data from all three tasks. In this plot, as in all subsequent plots with similar conventions, the distinction between different tasks is retained at the level of the marginal histograms (plotted to the sides of the co-modulation plot; please refer to caption for details). As detailed in Appendix A, human datasets from the other two tasks (polarity discrimination and energy extraction) are less structured than the dataset from the detection task (**Figure 5B**). Notwithstanding such differences, the main conclusion prompted by the detection dataset is applicable to those additional tasks: when assessed with reference to the commonly adopted metric of % correct responses, Lenet-5 appears to capture human behaviour; once the characterization is extended beyond this metric, glaring discrepancies become apparent.

### 3.3. Impact of different training specifications

We studied a wide range of variants around the training protocol detailed above, without much impact on our general observations. In the simulations discussed so far, Lenet-5 is trained on a given task (e.g. detection) and tested on that same task. We also attempted protocols in which the network was trained on different tasks and/or multiple tasks (Appendix B). **Figure 5C** shows the co-modulation plot associated with training the network on all three tasks simultaneously. In a variant of this protocol, we applied a different colour to the vertical markers placed above and below each stimulus array (red lines in **Figure 2**) to provide an explicit task-label (red for detection, green for polarity discrimination, and blue for energy extraction), but this modification had virtually no impact on network behaviour. With multi-task training, we notice a slight improvement over the single-training protocol in relation to the primary metric for zeroth- and first-order characterizations. For this reason, in the remaining part of this study we focus on networks that are primarily trained on all three tasks simultaneously. Besides the above-noted improvement afforded by this approach, it is desirable from a theoretical standpoint: ultimately, we would like to secure an account of human behaviour that is as comprehensive as possible within a framework that is as unified as possible. In additional exploration with different protocol variants, we also studied the role of stimulus characteristics at training (mixture of SNR/contrast levels and/or presence of external noise). As detailed in Appendix C, these variants do not substantially impact scatter across the co-modulation plot.

### 3.4. Transfer-learning and fine-tuning

The above results demonstrate that Lenet-5 does not develop human-like perceptual representations and/or computational motifs when specifically trained to maximize signal detection/discrimination for the same stimuli/tasks that were adopted with human observers. It may be argued, however, that this approach to training is inappropriate as a model of human learning: the human visual system is not entirely plastic; it is therefore unlikely that it can adjust its parameterization to the pervasive degree that is implemented by full training of a neural network [40]. Even if we accept that some form of back-propagation may be operating in biological substrates during learning [41], the human visual system would not afford this process unfettered access to every neuronal connection throughout its architecture from early visual cortex to decision-making structures, and for good reason: the human observers who participated in our experiments must carry out all sorts of visual tasks before and after collecting data for our study. It would make no sense that they re-structure large chunks of their visual system to suit our very specific and limited task requirements. They may re-structure small modules, and those may be larger than Lenet-5; even so, access to the modules and their final impact on behaviour would presumably be constrained by established circuitry that does not undergo substantial modification.

We can begin to address the issues discussed above by relying on a version of Lenet-5 that has been pre-trained to perform more ecological tasks, like natural scene classification. We then tweak this pre-trained model to perform our detection/discrimination tasks [42], and study whether the structure imposed by the ecological task steers the network towards developing properties that are more similar to those measured from humans. **Figure 6A** shows the co-modulation plot produced by fine-tuning Lenet-5 after pre-training on CIFAR-10: there is no improvement over training the network from scratch; in fact, the match with humans is either equivalent or poorer (compare **Figure 6A** with **Figure 5C**). This result suggests that the two training diets converge on similar model parameterizations. In other words, the effect of pre-training on CIFAR-10 may be largely washed out by fine-tuning.

**Figure 6.**
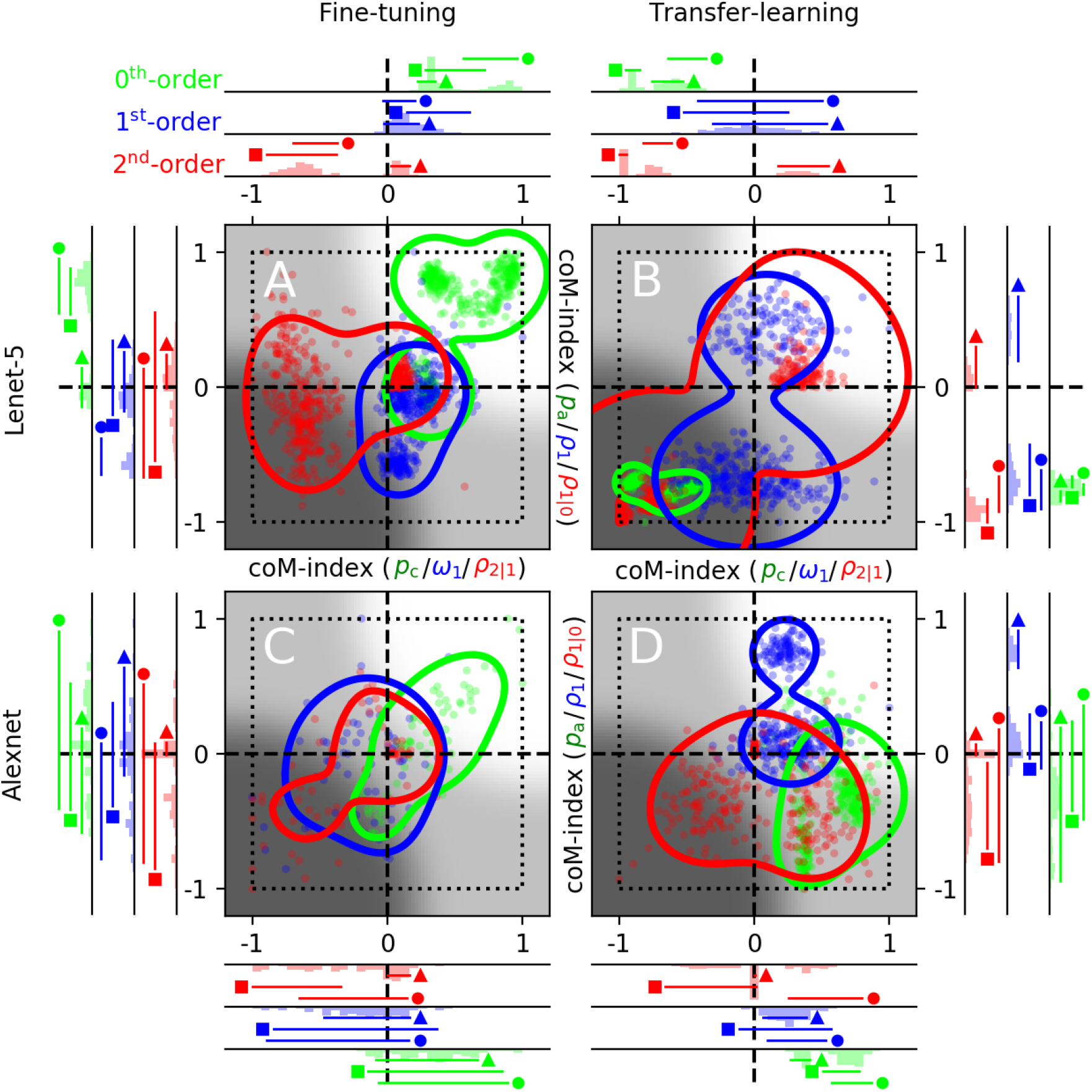
Fine-tunings/transfer-learning from scene classification tasks. Plotted to the conventions of **Figure 5. A**/**B** refer to Lenet-5 pre-trained on CIFAR-10 and subsequently fine-tuned/transfer-learnt to the detection/discrimination tasks used here. **C**-**D** show equivalent results for Alexnet pre-trained on Imagenet.

To enforce the pre-trained structure more effectively, we apply transfer-learning (only output layer is re-trained). Under this protocol, the network is almost entirely operating as specified by the scene classification task; the associated parameterization is essentially co-opted by the output layer to perform the detection/discrimination tasks. **Figure 6B** shows the resulting co-modulation plot: the network behaves in an entirely different manner than humans, despite learning all three tasks reasonably well (performance reaches 90% at the end of the training phase in **Figure 6A**) and performing above chance during the test phase (**Figure 6B**).

### 3.5. Potential role of network depth/architecture

All simulations detailed above refer to Lenet-5. As pointed out earlier, there is no reason to surmise that Lenet-5 may be too small to span the behavioural repertoire required by the problem at hand: this network can easily learn all three tasks, even when mixed within the same training protocol. Furthermore, it is able to produce higher-order descriptors that encompass the entire spectrum observed in humans. For example, the associated first-order descriptors may be lowpass (**Figure 4G**) or highpass (**Figure 7C**), and second-order descriptors may contain modulations that are relatively larger or smaller in amplitude than corresponding first-order modulations (see wide range spanned by radius variation for red circles in **Figure 4H** and **Figure 7D**). For the above reasons, we do not expect larger/deeper networks to outperform Lenet-5 in its ability to capture idiosyncratic human behaviour. Nevertheless, we explore this issue below to determine whether it may provide further insight into the relationship between network architecture and co-modulation index [43].

**Figure 7.**
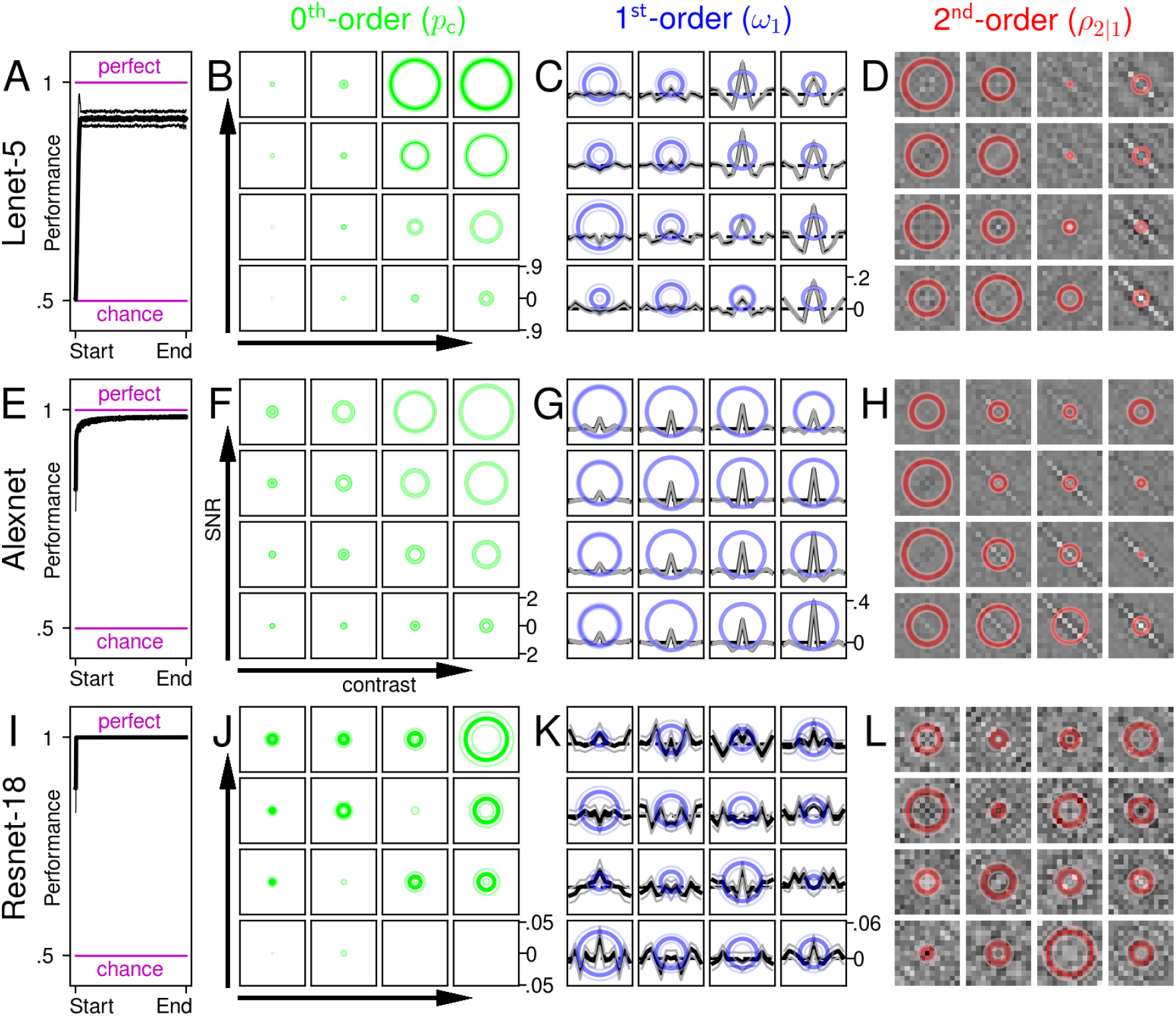
Transfer-learning deeper models does not improve co-modulation, and may even reduce it. **A**-**D** are plotted to the conventions of **Figure 4E-H**; here Lenet-5 is transfer-learnt to the detection task from pre-training on CIFAR-10. **E-H** show equivalent results for Alexnet, **I-L** for Resnet-18, both pre-trained on Imagenet. Although deeper networks learn more quickly during training (compare the progression from **A** to **E** to **I**), they do not produce more human-like behaviour during testing (compare **B-D** with **J-L** in relation to the corresponding human data in **Figure 4B-D**).

The co-modulation plots in **Figure 6C-D** were obtained by fine-tuning/transfer-learning Alexnet. When comparing these results with equivalent data from Lenet-5 (**Figure 6A-B**), there is no evidence that the larger architecture supported by Alexnet confers any improvement. We find similar results with other networks, whether smaller or larger than Lenet-5. For example, we constructed a reduced version of Lenet-5 consisting of only 3 layers, which we term Mini-3 (≈2K trainable parameters). This network acts directly on the 1-D vector defining stimulus intensity at each bar location and consists of a small convolutional layer (kernel size 13), the output of which is replicated through two branches: a simple-cell branch along which it is subjected to linear half-wave rectification, and a complex-like branch along which it is subjected to squaring. Outputs from the two branches are concatenated and submitted to a fully-connected layer that returns a binary label. Co-modulation indices for this network are comparable to those returned by Lenet-5.

As for larger networks, Resnet-18 produces extremely poor results. **Figure 7** plots full metric characterizations for the detection task in 3 different transfer-learnt networks: Lenet-5 (**A-D**), Alexnet (**E-H**) and Resnet-18 (**I-L**). In terms of achieving performance during learning, deeper is better: Lenet-5 cannot achieve perfect performance (**A**), Alexnet nearly achieves it (**E**), and Resnet-18 easily achieves it (**I**). However, the patterns produced at testing go in the opposite direction: Lenet-5 (**B-D**) is better aligned with the human trend (**Figure 4B-D**) than Resnet-18 (**J-L**).

We summarize the impact of network depth in **Figure 8A**. In this co-modulation plot, results from networks of increasing depth are overlayed as contours of increasing thickness/opacity. Co-modulation indices are *not* re-scaled separately for different networks/orders so as to make them directly comparable. There are two obvious features to this plot. First, only zeroth-order indices fall into the ‘good’ (bright) region, in line with the general finding discussed in previous sections. Second, co-modulation indices for the zeroth-order characterization follow a trajectory (indicated by orange line) that starts within the bright region for shallow networks, and progressively makes its way towards the origin of the plot for deeper architectures.

**Figure 8.**
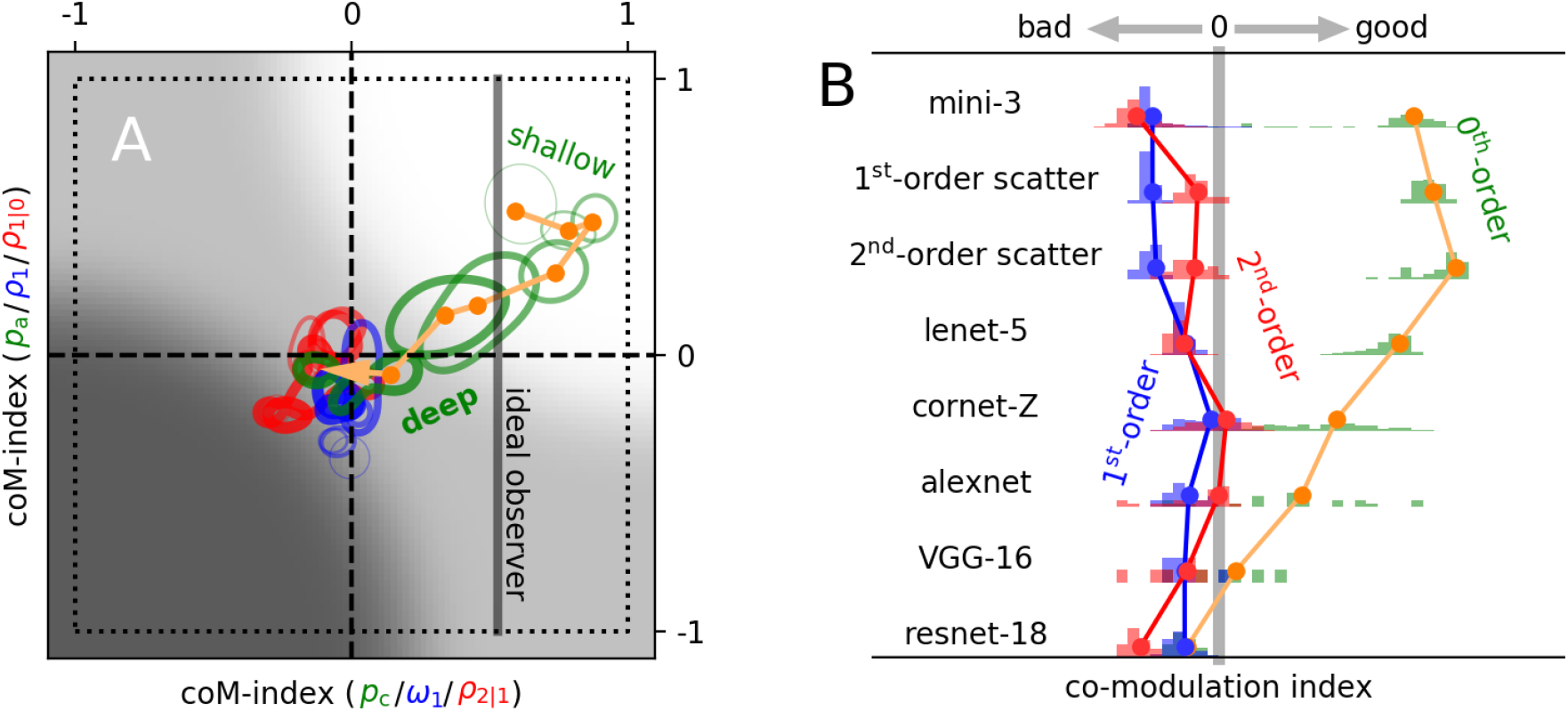
Network depth does not help. The co-modulation plot in **A** combines results from 8 different networks, each indicated by 3 different green/blue/red contours of increasing thickness/opacity with increasing network depth (dots for individual simulations are not plotted to avoid clutter). The orange trajectory proceeds from shallow to deep networks (see ordered list in **B**), demonstrating a reduction of the zeroth-order co-modulation index. Indices have *not* been re-scaled separately for different networks/orders to allow direct comparison. Black vertical line indicates co-modulation index associated with *p*_c_ values produced by the ideal observer (section 3.6). The x axis in **B** re-plots data from **A** after projection of each order onto positive diagonal of **A**. Histograms show distribution across different simulations for each order and each network separately.

**Figure 8B** plots data from **Figure 8A** using a different format, meant to further emphasize the effects outlined above. In this plot, the co-modulation index on the x axis is the average of the co-modulation indices from x and y axes in **Figure 8A**. Shallow networks, like mini-3 detailed above, achieve good zeroth-order co-modulation values (green histograms). The best results are returned by a scattering model parameterized for CIFAR-10 classification [19] (see caption to **Figure 8**). It may appear that there is no structure to first-order and second-order characterizations, although it is noticeable that for some networks (like mini-3) the corresponding indices are distributed clearly away from 0 towards negative values (see top blue/red histograms in **Figure 8B**). As we demonstrate further below, there are in fact important trends associated with these descriptors and their relationship to the zeroth-order characterization, as well as to other relevant metrics.

### 3.6. Theoretical bound on attainable performance

One of the many advantages associated with the stimulus/task specification adopted in this study is that it is possible to formulate an explicit ideal strategy for maximizing performance. This strategy is prescribed by ideal observer theory [26], and it simply amounts to template-matching against the target bar for detection and discrimination tasks (for the energy extraction task, the ideal strategy involves adding the exponentiated outputs of the bright-bar and dark-bar target templates [12]). The performance (% correct responses) associated with this strategy represents an upper bound on any model.

The ideal observer must be intended as a normative tool [45], not a biologically plausible account of human perception. First, it is a fully deterministic model (*p*_a_=1 under all conditions), while humans present a large amount of internal noise [29]. Second, for both detection and discrimination tasks, this model returns a first-order descriptor that is a rescaled image of the signal-to-be-detected (zero everywhere except for the central element) and featureless second-order descriptors, across all contrast/SNR conditions. This is patently inconsistent with the human measurements (**Figure 4B-D**). Nevertheless, it is a useful construct for understanding some limitations associated with the assessment of the potential match between more plausible models and human behaviour.

The black vertical line in **Figure 8A** marks the co-modulation index achieved by the ideal observer model for *p*_c_ (it is therefore a zeroth-order descriptor). It is not meaningful to compute a similar index for *p*_a_ because, as explained above, the notion of internal noise is not applicable to the ideal observer. It is noticeable that this ideal index is positive, indicating some degree of co-modulation between ideal performance and human performance. This is expected, because ideal performance depends on stimulus SNR, and it does so in a manner not too dissimilar than the human trend. However, ideal performance does not depend on contrast: it is for this reason that the ideal index falls below its counterparts from shallow networks (green outlines and orange data points lying to the right of the solid vertical line).

### 3.7. Theoretical bounds on attainable trial-by-trial prediction

From the above we can conclude that the analysis proposed in this study, namely the adoption of multi-order metrics with associated co-modulation indices, is certainly capable of recognizing that the ideal observer is inadequate for the purpose of capturing human behaviour. However, this is not the approach that is typically adopted in the literature. A more common approach is to rely on the trial-by-trial agreement between human label and network label, i.e. the proportion of trials on which human and network produce the same label in response to the same set of input stimuli [9]. A priori, this metric appears as a sensible choice for assessing human-network match on a fine scale. Yet we demonstrate below that, in fact, it is nearly useless for our application.

First, in order to evaluate human-model *p*_a_ on a meaningful scale, we need to set theoretical bounds on this quantity: what is the best achievable trial-by-trial match with the human response that any model can attain? Clearly, it is not possible to predict the human response on every trial with 100% accuracy because, as we have discussed above, humans do not behave in a deterministic fashion. In other words, even if we had access to a perfect model of the human brain, we would still fail at predicting human responses on some trials due to intrinsic variability of the human process [27, 29]. The upper limit on human-model 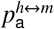 can be established via human-human 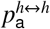: it is theoretically expected to lie within the interval 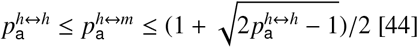 [44].

The upper/lower bounds predicted by theory are indicated by black/red solid lines in **Figure 9A**; 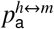 values associated with the scattering network model (y axis) fall within this range across SNR/contrast conditions (dot size reflects SNR, dot brightness reflects contrast). At face value, this result may appear encouraging and at least partly consistent with our earlier assessment that the scattering model produces markedly positive co-modulation indices for the zeroth-order characterization (green histograms in **Figure 8B**). However, 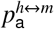 actually carries little power for the purpose of distinguishing between good and bad models (see below).

**Figure 9.**
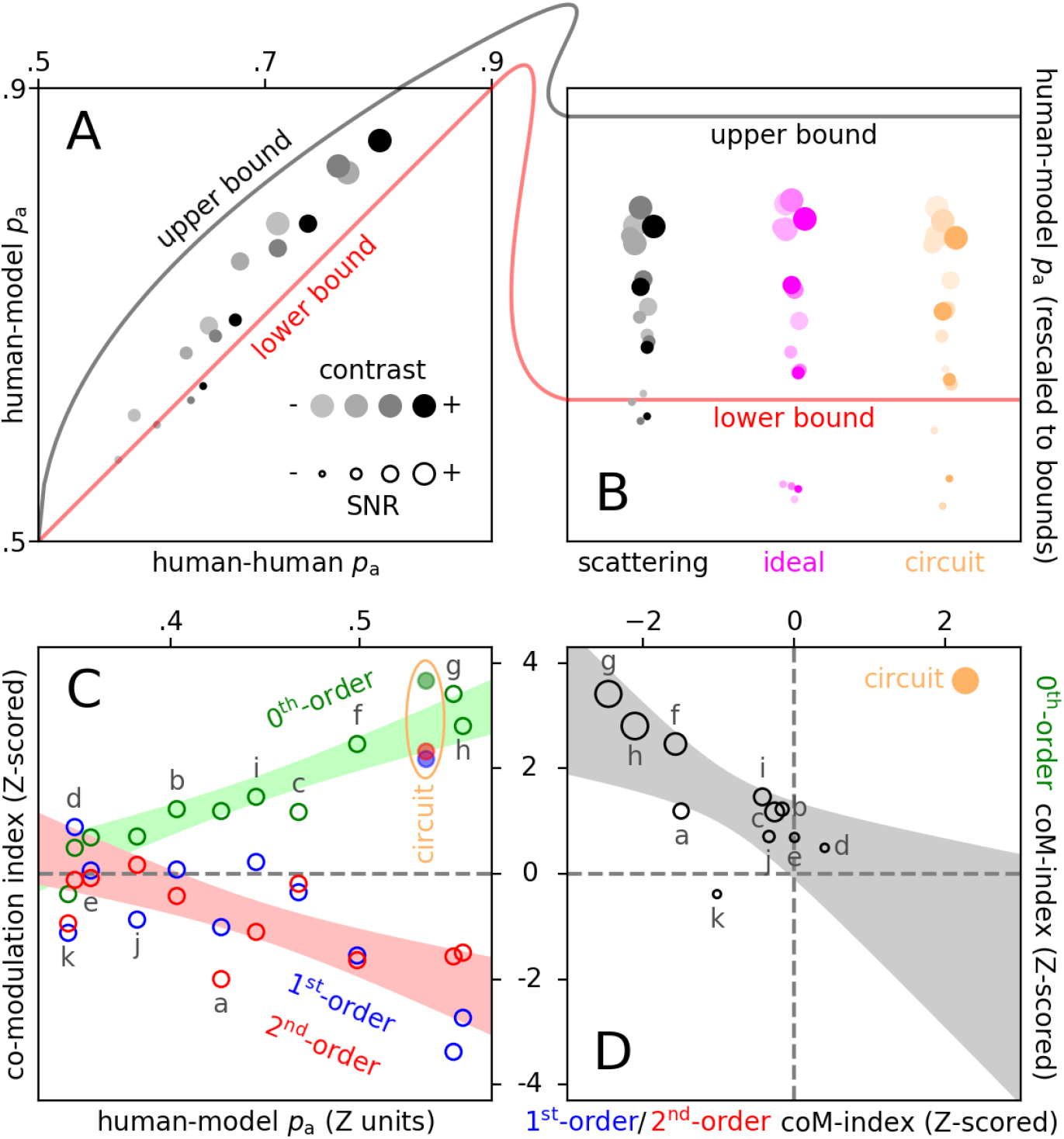
Human-model trial-by-trial agreement scales positively with zeroth-order co-modulation index, but negatively with higher orders. **A** plots human-model agreement (y axis) versus human-human agreement (x axis) for the 2-order scattering model [19] in the detection task across all 16 stimulus configurations (one dot per configuration; dot size scales with SNR, dot saturation scales with contrast). The theoretical limit on the former quantity is dictated by the latter and must lie within the region bounded by red/black lines [44]. Ordinate values from **A** are re-plotted by the black/gray symbols in **B**, after rescaling each human-model agreement value so that the corresponding lower and upper bounds map to 0 and 1. Equivalent data is plotted for the ideal observer (magenta) and the circuit model from **Figure 10B** (orange). Open symbols in **C** plot human-model agreement on x axis against Z-scored (mean divided by SD across simulations) co-modulation indices on y axis for 11 different networks/training-regimes: mini-3 (a); Lenet-5 trained on all tasks (b) or only detection (c), energy-extraction (d), polarity-discrimination (e); fine-tuned Lenet-5 (f); 1^st^-order scattering (g); 2^nd^-order scattering (h); fine-tuned CORnet-Z (i); fine-tuned Alexnet (j); fine-tuned VGG-16 (k). Solid symbols refer to circuit model (**Figure 10B**). Different colours refer to different orders: green for zeroth-order, blue for first-order, red for second-order. Green shaded area shows ±95% confidence intervals for linear fit of zeroth-order co-modulation index against human-model *p*_a_; the Pearson correlation coefficient for this fit is 0.94 (p<10^−4^). Red shaded area shows same for linear fit of average between first-order and second-order indices (coefficient of -0.79, p<10^−2^). Data from **C** is re-plotted in **D** with zeroth-order co-modulation indices on the y axis, average of first-order and second-order indices on the x axis, and symbol size scaling with human-model agreement. Shaded area shows linear fit (coefficient of -0.74, p<10^−2^) across all networks excluding circuit model (indicated by solid orange symbol).

**Figure 9B** plots 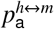 values (y axis) after rescaling each value so that the projected scale is anchored by corresponding upper/lower bounds; this plotting convention makes it easier to compare human-model agreement across conditions/models. The scattering data is plotted alongside corresponding data from the ideal observer (magenta): clearly, the latter is capable of achieving a degree of trial-by-trial agreement with the human responses that is entirely comparable with the values attained by the scattering model. Based on human-model agreement alone, we would conclude that the ideal observer is just as adequate as the network model associated with largest zeroth-order comodulation index. On the contrary, we know from the in-depth analysis adopted in this study that the ideal observer is *not* capable of capturing numerous features of the human process (section 3.6) and presents substantially lower *p*_c_ co-modulation than the scattering model (**Figure 7A**). In other words, human-model agreement is a poor metric for model selection.

Next we ask how human-model agreement relates to the multi-order characterization introduced previously. **Figure 9C** plots Z-scored co-modulation indices (mean divided by SD across simulations) for a range of networks and training protocols on the y axis, versus corresponding human-model agreement (x axis). These two quantities are clearly related, and in opposite directions for different orders: the zeroth-order co-modulation index scales positively with agreement (green open circles), while first-order/second-order co-modulation indices scale negatively (blue/red open circles). In other words, as networks acquire better trial-by-trial agreement with the human response, they also produce more human-like trends for performance and internal noise; however, they also develop opposite-to-human trends for higher-order descriptors (see caption to **Figure 9** for statistical assessment).

We conclude this section by relating our metric of preference, namely bound-normalized 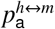 (plotted on the y axis in **Figure 9B**), to other existing measures of similar quantities such as Cohen’s κ, designed to factor out the inevitable co-variation between 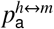 and *p*_c_ of both human and model [46]. In SDT-based approaches, this quantity is factored out by the SDT model itself that is utilized to estimate internal noise intensity [28, 30, 29]. In probabilistic approaches, it is factored out by computing a baseline reference under the assumption of independence between human and model [47, 9]. In our approach, it is factored out indirectly via 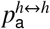, as this quantity also scales with *p*_c_. These different approaches are clearly inter-related. For example, the correlation coefficient between our normalized metric and corresponding Cohen’s κ values across models and conditions is 0.9 (p< 10^−8^). Our observations therefore apply to these other metrics too. We favour normalized 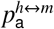 because it also carries information about theoretical bound saturation for optimal prediction [44], a quantity that is not accounted for by the above-mentioned approaches.

### 3.8. Tailored circuit model

It seems legitimate to wonder what kind of network, if any, may reproduce human behaviour. In particular, what size would be required for such a model, and what degree of complexity? Based on the observation that network depth may not be beneficial and may even be detrimental (section 3.5), we expect that a relatively simple circuit should be sufficient to provide a satisfactory account. Furthermore, because the cognitive phenomenon of interest has been extensively studied over decades of vision research [48, 49], we expect that known computational motifs should be involved, without the need to invoke fanciful models. Indeed, as we have demonstrated in prior work, a simple circuit can account for the human data [12]. **Figure 10B** reproduces the structure of this circuit, and **Figure 10A** projects its output in the detection task onto co-modulation space for direct comparison with the other models we have considered so far. Clearly, this simple model outperforms all other models, in that it captures human behaviour across all three orders of characterization.

**Figure 10.**
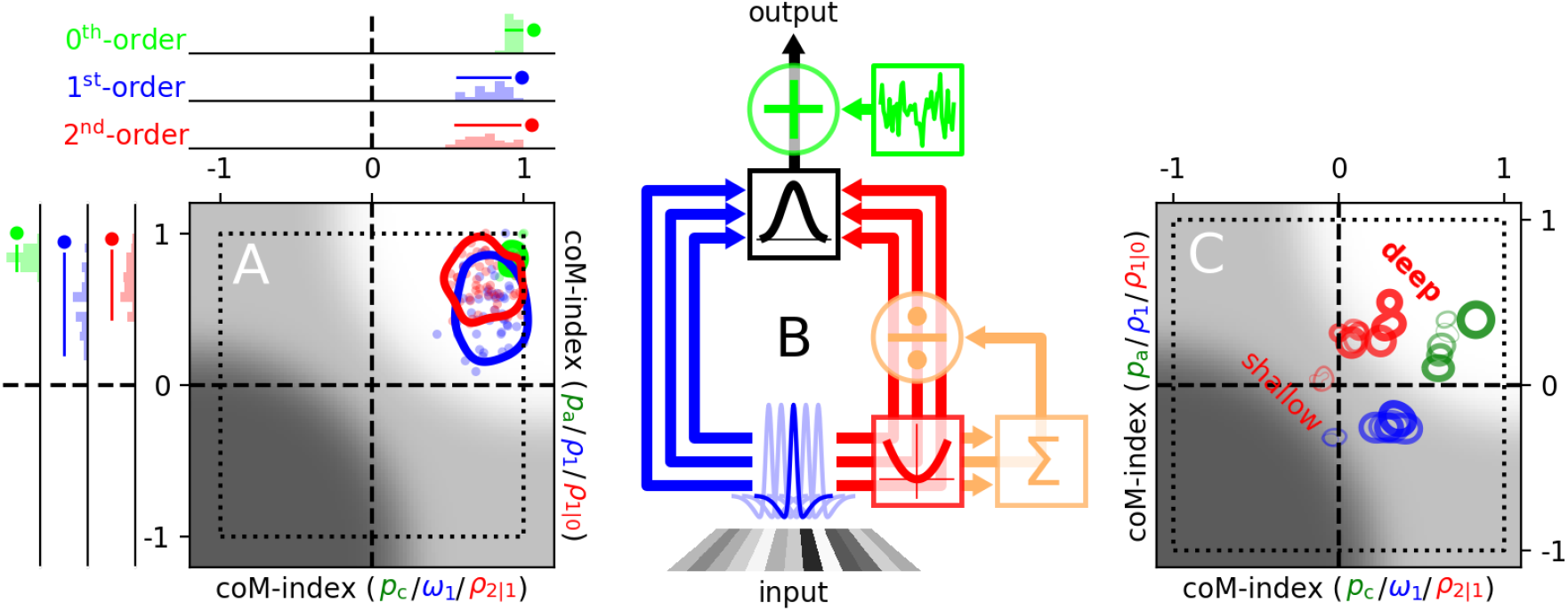
Circuit model captures human detection dataset and supports training of larger networks. **B** summarizes the architecture of the circuit. First, 1-D input stimuli are convolved with a bandpass filter (blue Mexican-hat shaped profiles). Outputs from the filter bank are then routed through two branches: a linear branch (left, blue) along which they are left unmodified; a nonlinear branch (right, red) along which they are subjected to squaring (red square element) and divisive gain control (orange elements). The two resulting outputs are Gaussian-weighted (black square element at the top), summed and corrupted by a fixed additive noise source (green element). Please refer to [12] for detailed implementation of this model. The associated co-modulation plot (**A**) produces excellent results across all three orders of characterization (all data points fall within bright region). This model can be used to produce binary labels for a large set of noise-only stimuli; the resulting labelled dataset can then be fed to deep networks for training. **C** shows progression across the co-modulation plot for 8 networks trained in this manner: Lenet-5, 2^nd^-order scattering, CORnet-Z, Alexnet, VGG16 and Resnet-18 (all fine-tuned except scattering). Increasing depth (in the order just listed) is indicated by thickness/opacity of contours.

In order to study more closely how the circuit model compares to the trained network models, its co-modulation indices are plotted in **Figure 9C** using solid symbols, against its human-model agreement. It is apparent that this model falls within the high-performing range for zeroth-order alongside the scattering models (indicated by letters g and h), without loss of power for higher orders (blue/red solid symbols fall above the horizontal dashed line).

**Figure 9D** re-plots data from **Figure 9C** using a different format, intended to emphasize just how starkly the circuit model departs from the operating characteristic spanned by the other network models. In this panel, zeroth-order comodulation indices are directly transferred from the y axis in **Figure 9C** to the y axis in **Figure 9D**; as for the x axis in **Figure 9D**, it plots the average of first-order and second-order co-modulation indices from the y coordinates of blue/red data points in **Figure 9C**. Symbol size scales with human-model agreement (x values from **Figure 9C**). The inverse relationship between zeroth-order indices and higher-order (first-order+second-order) indices is now apparent: whatever is gained by a given network model along the zeroth-order axis (y), it is concomitantly lost along the higher-oder axis (x). In stark exception to this trend, the circuit model (solid orange symbol) falls within a different quadrant (upper-right).

### 3.9. Tailored circuit as proxy for the human process during training

In an effort to determine whether neural networks can be made to resemble human behaviour under any circumstance, we have made numerous attempts at pushing convergence towards the human pattern by training networks against the human label, rather than the signal label: instead of supervising via the correct label specified by the task (correct/incorrect), we apply supervision with reference to the binary response produced by humans (the training set consists of the same stimuli used in the psychophysical experiments). Our attempts were unsuccessful regardless of architecture, whether we train on one task or all tasks, whether we train on all trials or only on those for which humans produced an incorrect response (under the rationale that incorrect trials may carry more information about the idiosyncracies of human behaviour), and other variants attempted over several pilot simulations. These results may appear to indicate that neural networks cannot represent the human process even when pushed to do so, however we must first entertain a simpler explanation: the human dataset is inevitably limited in size and, more importantly, it is very noisy due to intrinsic variability (human-human agreement <1). It is therefore possible that it does not carry sufficient statistical power to drive networks towards the configuration that represents the human process.

To overcome the above limitations of the human dataset, we utilize the circuit-model as proxy for the human process. As demonstrated in section 3.8, the circuit-model provides an excellent match for the human process in the detection task, and its output carries human-like information across all three orders of characterization (**Figure 10A**). For the purpose of generating a training dataset, this model presents two advantages over the human process: first, we can create a potentially unlimited number of labelled trials; second, we can turn off internal noise within the model (green element in **Figure 10**), so that the resulting simulated responses are deterministically coupled to the stimulus and statistically robust. In our first series of attempts, we trained networks in the detection task against labels produced by the circuit-model for a large training set of noisy mixed-SNR/contrast stimuli (>1 million), often assembled into sizeable mini-batches; these attempts were unsuccessful in the sense that, although some networks did learn zeroth-order human structure, they failed to reproduce higher orders.

In evaluating the disappointing results detailed above, we consider that, particularly for mid-to-high SNR trials, responses from the circuit-model are predominantly driven by the target signal and only mildly modulated by the external noise process. First-order/second-order descriptors, however, are precisely computed by leveraging on those smaller modulations that are noise-coupled (section 2.3). We therefore attempt a training protocol with input stimuli containing only noise; the corresponding label is produced by the circuit-model in response to those noise samples. There is no correct/incorrect response for this training set, so there is a legitimate concern that networks may not learn the detection task at all: they are never explicitly given any feedback regarding the signal detection task itself. However, if networks learn the structure of the circuit-model from the input-output coupling between the noisy process and the associated label, they may be expected to perform the detection task successfully when target signals are reintroduced at testing.

Indeed, this is what we find: not only several networks are able to perform detection successfully during the testing phase, but the associated first-order/second-order characterizations are much better aligned with the human process than under any other protocol we have attempted in this study. For example, Resnet-18 now captures the human transition from bandpass to lowpass with contrast (**Figure 11K**, compare with **Figure 4C**); it also reproduces the larger amplitude of second-order descriptors at low contrast/SNR (**Figure 11L**, compare with **Figure 4D**). By learning these two orders, the network also learns an excellent zeroth-order rendition of the human pattern (**Figure 11J**, compare with **Figure 4B**). We find that this result is obtained with a smaller number of training samples when fine-tuning from a pre-trained parameterizations via Imagenet, however the same result is obtained when training from scractch as long as a sufficient number of training samples is adopted. In other words, pre-training on a classification task is helpful, but not necessary.

**Figure 11.**
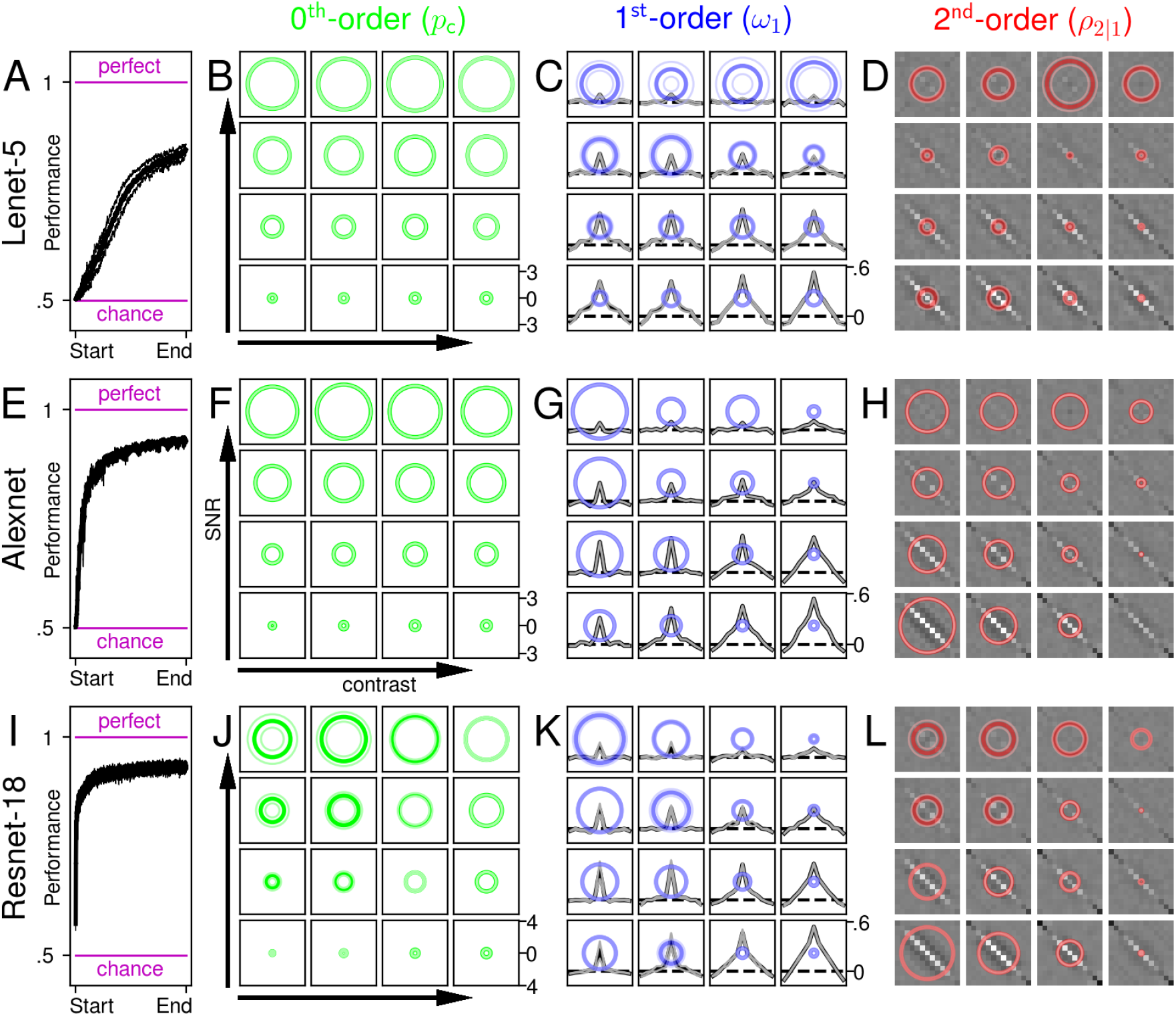
Deeper models are capable of learning labels produced by the circuit-model in response to noise-only stimuli. Plotted to the conventions of **Figure 7**. Networks are trained with input stimuli containing no signal-to-be-detected, only noise. Correct/incorrect labels are replaced with labels generated by the tailored circuit model (**Figure 10B**). Under these conditions, Resnet-18 produces descriptors that match human counterparts across all orders (compare **J-L** with **Figure 4B-D**).

For this protocol, we also find positive dependence on network depth: Lenet-5 is slow at learning the circuit-model and reaches plateau at about 70% successful reproduction of associated labels during training (**Figure 11A**), while also failing at capturing the contrast-dependence of all three orders of characterization (**Figure 11B-Figure 11D**). Alexnet learns the circuit-label more quickly/efficiently (**Figure 11E**) and captures the bandpass-to-lowpass first-order transition with contrast (**Figure 11G**), however it does not reproduce the contrast-dependence of the zeroth-order metric (**Figure 11F**). Resnet-18 improves on all of the above (**Figure 11I-L**). This state of affairs is summarized in **Figure 10C**: deeper networks move into the bright region of the co-modulation plot for both zeroth-order (green contours) and second-order indices (red contours); for the first-order index (blue contours), although all networks fail to capture the complementary metric (ρ_1_ on the y axis), they do capture co-modulation for the primary metric (ω_1_ on the x axis). These results go in the opposite direction to those obtained without guidance from the circuit-model (compare **Figure 11** with **Figure 7**).

## 4. Discussion

### 4.1. Strengths and weaknesses of the proposed approach

This study capitalizes on two essential design elements: extreme simplicity of input specifications, and extreme complexity of output characterization (**Figure 1**). Both come with advantages and disadvantages.

The advantages associated with simplified stimulus/task protocols are numerous: 1) they allow quantitative specification of stimulus manipulations such as SNR/contrast changes; 2) they greatly reduce ambiguity in the interpretation of specific effects, for example by largely mitigating concerns about levelling the playing field for human-machine comparison [2, 3]; 3) they support ideal observer analysis [34], often unavailable for more complex tasks like object classification; 4) they align with theoretical results, such as maximum bounds on point (i.e. trial-by-trial) prediction [44]; 5) the associated neural network representation is not expected to involve articulated features such as recurrency as required by more complex tasks [50]. The most obvious disadvantage is that it is unclear how these results may relate to more complex stimuli/tasks. In the context of this study, however, this is not a relevant limitation: the motivation here is to demonstrate that deep neural networks cannot be trained to account for human behaviour; a conclusion of this kind is stronger if demonstrated for simple tasks, because it is reasonable to assume that it may also apply to more complex tasks. When this logic is turned around, namely to claim that failure at a complex task should imply failure at simpler tasks, it becomes less compelling.

The primary advantage associated with a complex behavioural characterization is that it carries greater power for model selection: as we have demonstrated here (and discuss further below), the class of multi-layered descriptors afforded by this analysis imposes stringent constraints on candidate models of the human process, beyond what is afforded by mainstream metrics such as detection performance or human-model point agreement (sections 3.6-3.7). There are potential disadvantages, however. First, descriptors of the kind proposed here are only supported by certain stimuli, notably those that carry external noise: in the absence of an external perturbation source, it is not possible to derive first-order and second-order descriptors. Second, reliable characterization of said descriptors requires an unusually large data mass: the psychophysical dataset used in this study consists of > half-a-million human responses [12]. Third, there is a degree of arbitrariness in the choice of metrics, although those presented here are motivated by general theoretical considerations [15] and encompass all documented descriptors for stationary detection processes. Fourth, the co-modulation index is a construct that requires stimulus manipulations (like contrast/SNR here) that may be undefined and/or meaningless in certain applications.

Within the context delineated above (which we discuss further in section 4.4), the methodology adopted in this study has exposed important limitations of machine learning for the purpose of reproducing human behaviour, however it has also left numerous questions unanswered. In particular, it is unclear where the limitation lies specifically. We consider two possibilities: 1) architectural limitations, whereby neural networks do not carry enough computational power to represent the human process; 2) learning limitations, whereby networks are in principle capable of representing the human process, however they are not capable of converging to such representation via learning under conditions comparable to those adoped with humans. We discuss both of them in more detail below.

### 4.2. Representational power of deep architectures

It is extremely unlikely that the bottleneck may be represented by insufficient circuitry: we have demonstrated that the human pattern can be reproduced by a relatively simple structure consisting of no more than a convolutional layer, a static nonlinearity (squaring) and divisive gain control (**Figure 10**). It is inconceivable that deep networks such as Alexnet cannot, at least in principle, be parameterized to represent this level of computation. To demonstrate this explicitly, we have identified a specific training protocol that allows networks of reasonable size (CORnet-Z and above) to represent the human process adequately (**Figure 10C**); the key ingredient of this protocol is the removal of target signal from the training stimulus set (section 3.9). To understand how this modification may render networks capable of learning higher-order structure, we discuss below a number of connected issues.

Consider the relationship between zeroth-order and first-order characterizations. In its simplest form, zeroth-order is intended here as the proportion of correct responses, while first-order refers to the first-order kernel derived from noise [30]. These two descriptors are only loosely connected, in the sense that the same first-order kernel may be associated with different performance values, and the same performance value may be associated with different first-order kernels [16]. Similar considerations apply to the potential connection between those orders and the secondorder descriptor. Because of this partial inter-order independence, there is a sense in which the different orders must be learnt separately: it is not the case that learning the structure associated with one order naturally carries along other orders; under some circumstances, learning the zeroth-order human structure may even go hand-in-hand with anti-learning higher-order characteristics (section 3.7), where by ‘anti-learning’ we mean learning patterns that are *opposite* to those displayed by humans (**Figure 9C-D**).

With the above in mind, we can begin to understand the most likely source of benefit afforded by removing the target signal: zeroth-order information is essentially unavailable in the absence of such signal. The zeroth-order metric is intimately connected with percentage of correct target identifications: in the absence of the target signal, this quantity remains undefined. By removing this signal, the only learnable content is dominated by higher-order characteristics of the underlying input-output process, so neural networks can capitalize on those and learn them effectively. In the presence of the target signal, the training dataset is dominated by zeroth-order characteristics; network learning is therefore dominated by those, at the detriment of learning higher-order characteristics. Our results also indicate that, at least for the experimental conditions of this study, learning higher-order structure carries along human-like zeroth-order structure ‘for free’ (section 3.9).

### 4.3. Training like a human

Based on the above, we must conclude that the discrepancy between the structure learnt by neural networks and the corresponding human process derives from the training protocol. This issue sits at the very core of this study, because the primary attraction of applying neural networks to human data of the kind discussed here is that network models may provide a principled explanation for the specific structure exhibited by humans [51, 2, 3]. As a static structure (i.e. without an account of its developmental trajectory), there is no need for further explanation from deep networks: we already possess a suitable model in the form of the tailored circuit (section 3.8). This model approximates human behaviour in the detection task to second-order (**Figure 10A**), it is small (**Figure 10B**), and it is physiologically plausible [12]. However, it does not explain *why* humans present the particular structure it is designed to model. Neural networks become attractive in this specific context: if we can train them to display human-like behaviour after exposure to meaningful learning scenarios, this may provide important insights into why humans develop certain sensory properties.

What is a ‘meaningful’ training protocol? Within the context of this study, meaningful refers to carrying some non-trivial explanatory power. For example, the training protocol involving supervision by the tailored circuit is not meaningful. Although it involves some non-trivial design choices (such as removal of the target signal), the resulting networks do not present any explanatory power beyond the tailored circuit model: they merely reproduce that model (section 3.9). An example of meaningful protocol is the transfer-learning paradigm (section 3.4): the rationale behind this approach is to demonstrate that networks pre-trained on a higher-level task, such as object categorization, will develop specific structural properties that may be exposed by low-level tasks, such as detection/discrimination of simple visual elements as adopted in this study. A result of this kind would be conceptually interesting, and may offer ecological insight into the underlying mechanisms [42]. Unfortunately, as we have documented throughout this study, we were unable to identify a successful training protocol that would classify as meaninfgul.

### 4.4. Significance of higher-order characterization

Our failure to identify a non-trivial training protocol that leads to human-like architecture does not imply that such a protocol does not exist: it is very well possible, in fact almost certain, that such a protocol does exist. Regardless, our study exposes important weaknesses associated with the manner in which this problem is approached by mainstream investigations in machine learning: had our evaluation of the potential match between network and human behaviours been based solely on zeroth-order metrics, as is almost invariably the case in current literature, we would have concluded that neural networks reproduce human behaviour to an excellent degree. It should be noted that this conclusion would not have seemed trivial: although it is certainly trivial that human performance scales with stimulus SNR (a feature that is captured by the simplest conceivable model), it also varies with stimulus contrast, a characteristic that is non-trivial and unexpected from a normative standpoint (the ideal observer predicts no such trend). Similarly, double-pass agreement presents pecularities that are not straightforward consequences of simple model architectures [12].

The above observation applies regardless of the possibility that an appropriate training paradigm, combined with a suitable network, may reproduce human behaviour: such paradigm would not be sought in the absence of higherorder characterization, because it would seem unnecessary to do so. It is specifically the level of analysis afforded by first-order/second-order metrics that brings about the realization of the discrepancy between network and human behaviour. It is therefore imperative that we discuss the potential limitations associated with this analysis, which we have anticipated in section 4.1.

Because the application of external noise is central to the paradigm introduced here, we must consider how it generalizes to other stimuli/tasks. With relation to the statistical properties of the noise source, it should be noted that higher-order descriptors may be computed, at least in principle, for any class of external perturbations, not just Gaussian white noise as adopted in this study. Clearly, white noise presents some attractive properties that are well understood and is therefore desirable under many experimental settings [52], but perturbation-based descriptors can be derived from biased and highly structured stimuli, such as those associated with natural statistics [53].

A more pressing issue is whether results obtained with noisy stimuli are relevant to noiseless conditions. This question is not answerable in general terms: we cannot make a statement about all possible stimuli and tasks. However, it is answerable in relation to the specific stimuli and tasks used here: a prior study has reported descriptors analogous to the first-order class examined here, but obtained using a different technique (sub-threshold summation) that involves noiseless stimuli [54]. The effect of contrast measured by that study is virtually identical to the effect we examine here (bandpass-to-lowpass transition with increasing contrast) and is aligned with other relevant measurements [55], providing clear evidence that our results do transfer to conditions where stimuli are devoid of external noise.

A second important issue connected with higher-order estimation is the specific choice of scalar metrics. There are two separate issues here: the choice of descriptor, and the choice of derived metric. For example, the first-order descriptor is the full first-order kernel (each black trace in **Figure 4C**), while the derived metric is the spectral centroid of that descriptor (each blue circle). There are strong theoretical motivations behind the choice of descriptor: it is widely used in both electrophysiology [56] and psychophysics [14]; for a linear-nonlinear model (also termed template matcher), it provides a full characterization of the perceptual mechanism [30]; from the standpoint of descriptive statistics, it is essentially the only sensible choice for extracting first-order information from the coupling between external noise source and perceptual process [57]. Similar considerations apply to the second-order descriptor [15].

Contrary to the above, the coice of metric is largely arbitrary. At the same time, it is not mandatory: full firstorder/second-order descriptors are the relevant object of reference. In this study, we adopt specific metrics (such as spectral centroid) because they capture important aspects of the manner in which the first-order descriptor varies as a function of stimulus contrast/SNR [12]. This choice is therefore primarily dictated by the manipulations adopted in this study; in other contexts, different metrics may be more appropriate. What matters is that no such choice is necessary in order to implement the approach utilized in this study. Related considerations apply to the co-modulation index: being derivative of the scalar metrics, it is arbitrary and tailor-designed for application to the conditions of the experimental investigation examined here. However, it is not necessary to rely on this particular index to draw meaningful conclusions from higher-order characterizations of behaviour.

### 4.5. Integration with current knowledge on the application of DCN’s to human behaviour

Although different in many respects, the approach introduced here presents some points of contact with existing efforts in the literature. Previous psychophysical characterizations of deep networks have exposed important discrepancies between certain network architectures and human behaviour [58, 59]; in some instances, those discrepancies have been rectified via the addition of specific components, such as recurrent structures [50, 10]. An interesting example from this class of studies is represented by recent work on ‘uncrowding’, a perceptual phenomenon whereby the impact of flanker stimuli onto an embedded target is not easily predicted from flanker structure [59]. Feed-forward architectures cannot account for this phenomenon, however networks that incorporate grouping processes are able to provide a satisfactory account of the human data [10], in line with related work in neurophysiology [50]. In these studies, leverage into the comparison between humans and networks is achieved via more articulated stimuli rather than depth of behavioural characterization as implemented here. In some instances, perturbation-based techniques have been exploited to dissect network structure on a finer scale that bears superficial similarities to the first-order description adopted here [60], but in a very different context (face processing) for which most of the tools used here (e.g. ideal observer analysis) are not available.

A separate but related line of research has examined the relationship between human and network behaviour on a trial-by-trial basis [9], an approach known as ‘molecular’ psychophysics [27]. This approach shares similarities with (but also differs from) sample-by-sample comparisons of error production in humans versus networks [50]. As we have demonstrated in section 3.7, at least for the conditions of our experiments and tasks, little information is gained from this type of metric: it is incapable of discriminating between patently inaccurate and substantially accurate implementations of the human process, such as the ideal observer on the one hand, and the tailored circuit model on the other (**Figure 9B**). Furthermore, this class of metrics (which we term human-model agreement here) correlates positively with the zeroth-order characterization in our study, but not with higher-orders (**Figure 9C**).

We propose the following plausible explanation for the reduced resolving power of this metric when applied to the context of our experiments. In prior applications, the classifier (whether human or artificial) produces multi-label outputs for relatively higher-level tasks such as image classification [9, 50]; in these paradigms, heterogeneity of the output response is driven by *class* heterogeneity of the input (many different unperturbed image classes). Our design, on the other hand, involves binary labels for low-level signal detection/discrimination; in such protocols, response heterogeneity is driven by stimulus *perturbations* of just two basic image classes (few different images perturbed in many different ways). The importance of this distinction becomes clear when one considers that most of the analytical tools employed in our study are poorly transferable to image classification of noiseless natural scenes [9, 50]. From our viewpoint, this distinction points to the effectiveness of our approach in exposing limitations beyond those highlighted by prior work.

We emphasize that the purpose of this study is not to belittle deep convolutional networks, but rather to contribute towards ensuring that their full potential is unlocked adequately. As we have demonstrated in **Figure 11**, depth does play a role in conferring greater ability to learn the full human representation (albeit via intercession of the circuit model), however this conclusion can only be reached through inspection of higher-order descriptors. If the analysis is limited to the zeroth-order characterization, following routine practice in the evaluation of biological versus artificial behaviour, the opposite conclusion may be reached [43]: depth appears detrimental (**Figure 7**). We are confident that artificial networks carry the potential to explain the human pattern characterized here: after all, we know that a simple circuit is able to account for it (section 3.8). The critical question is *why* humans come to engage this specific mechanism for solving a task which, by itself, does not call for that particular mechanism (the ideal observer does not implement it). The answer to this question must come from incorporating knowledge beyond the task itself, possibly through learning other tasks (transfer learning) or via limitations of hardware implementation (in-built neural circuitry with limited plasticity). Either way, neural networks represent powerful tools for answering this question of paramount conceptual significance. To deliver on this potential, they must be equipped with suitable tools for assessing the structure of their output in relation to human behaviour. Our study offers some tentative pointers on how to push forward in this direction.

## Acknowledgments

Supported by the Agence Nationale de la Recherche (ANR-16-CE28-0016, ANR-19-CE28-0010-01, ANR-17-EURE-0017, ANR-10-LABX-0087 IEC, and ANR-10-IDEX-0001-02 PSL*).

---

**Figure .12.**
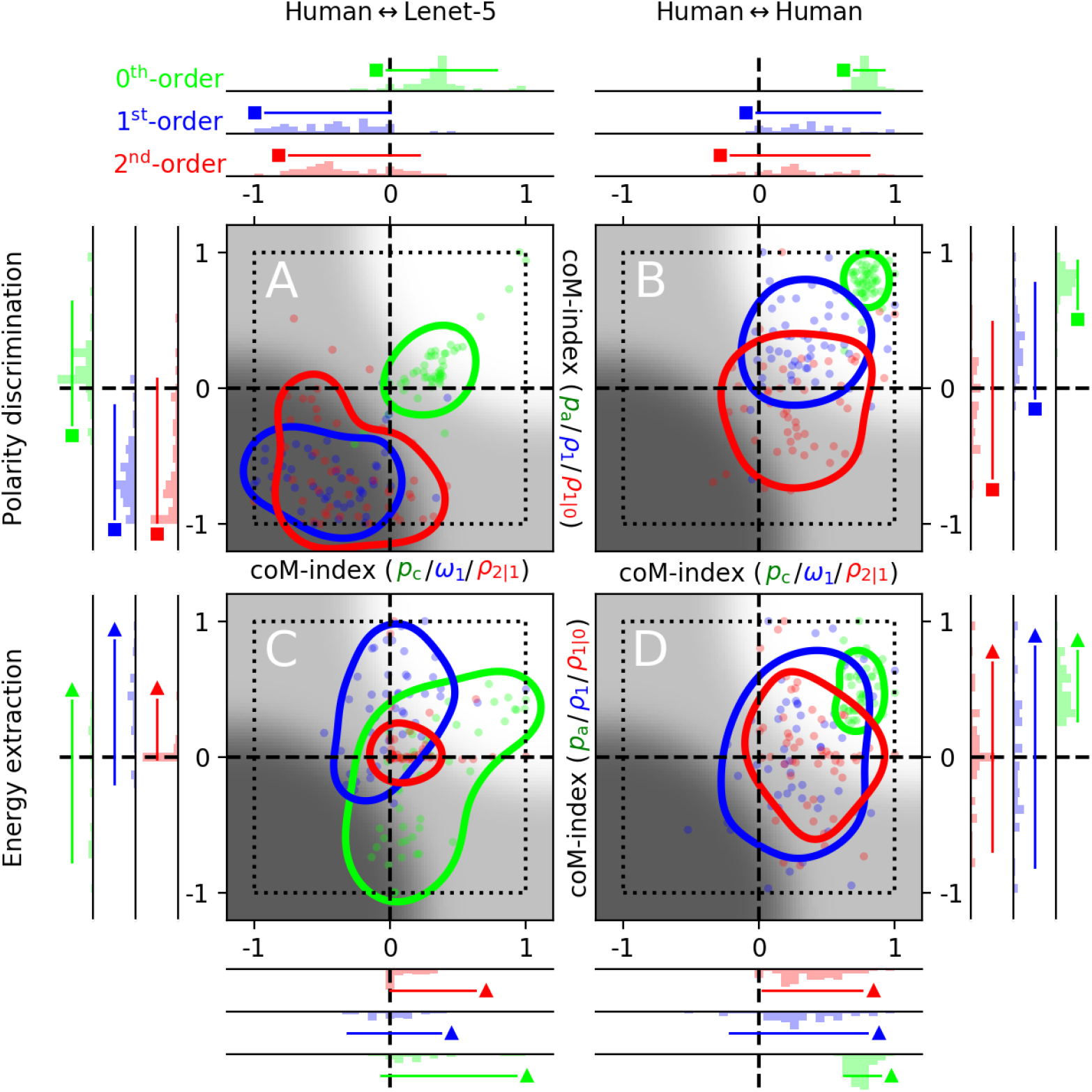
Human dataset is unevenly structured for different tasks. **A-B** are equivalent to **Figure 5A-B**, except they plot results for the polarity-discrimination task (**Figure 2D-F**), while **C-D** plot results for the energy-extraction task (**Figure 2G-L**).

## Appendix A

Polarity-discrimination and energy-extraction tasks

From the perspective of the extent to which Lenet-5 captures human behaviour, **Figure 5A** demonstrates a qualitative jump associated with the transition from zeroth-order to higher orders of behavioural characterization: at zeroth-order, the network does excellently; at higher orders (first-/second-), it fails miserably. An important question is whether this result is specific to the detection task (**Figure 2A-C**) or whether it generalizes to other tasks. **Figure.12A** shows the co-modulation plot associated with the polarity discrimination task (**Figure 2D-F**); similarly to the detection task, co-modulation is good for the zeroth-order characterization (green data falls within bright region) but poor for higher orders (blue/red data falls within dark region). More specifically, the first-order characterization for the network displays a dependence on contrast/SNR that is *opposite* to the human pattern (blue data falls within lower-left quadrant corresponding to negative co-modulation indices); the second-order characterization shows no co-variation between network and human trends (red data points tend to fall around the origin, with a mild tendency towards the lower-left quadrant). **Figure .12C** shows the co-modulation plot associated with the energy extraction task (**Figure 2G-L**); except for mild success in reproducing the human pattern for *p*_c_ (green data points largely fall to the right of the vertical dashed line), the network fails across the board.

Not all of the above failures are genuine. As pointed out in section 3.2, near-zero co-modulation may be an inevitable consequence of unstructured measurements from human observers. For this reason, it is necesssary to check that the human measurements do present structure by inspecting the corresponding human-vs-human comodulation plots in **Figure .12B,D**. For the polarity discrimination task, structure is present at both zeroth- and first-order (green/blue data points fall within upper-right quadrant); the second-order characterization, however, is only structured with respect to the primary metric (ρ_2|1_) plotted on the x axis (red data points fall to the right of vertical dashed line), and presents no structure for the the complementary metric (ρ_1|0_). This means that the apparent failure on the part of Lenet-5 to capture human trends with respect to this metric (y coordinates of red data points in **Figure.12A**) is a moot point. Similar considerations apply to the energy-extraction task: both first-order and second-order characterizations appear to lack structure with relation to the complementary metric (blue/red data points in **Figure.12D** scatter around the horizontal dashed line), invalidating the interpretability of the corresponding failures on the part of Lenet-5 (blue/red data points in **Figure .12C** also scatter around the horizontal dashed line).

**Figure .13.**
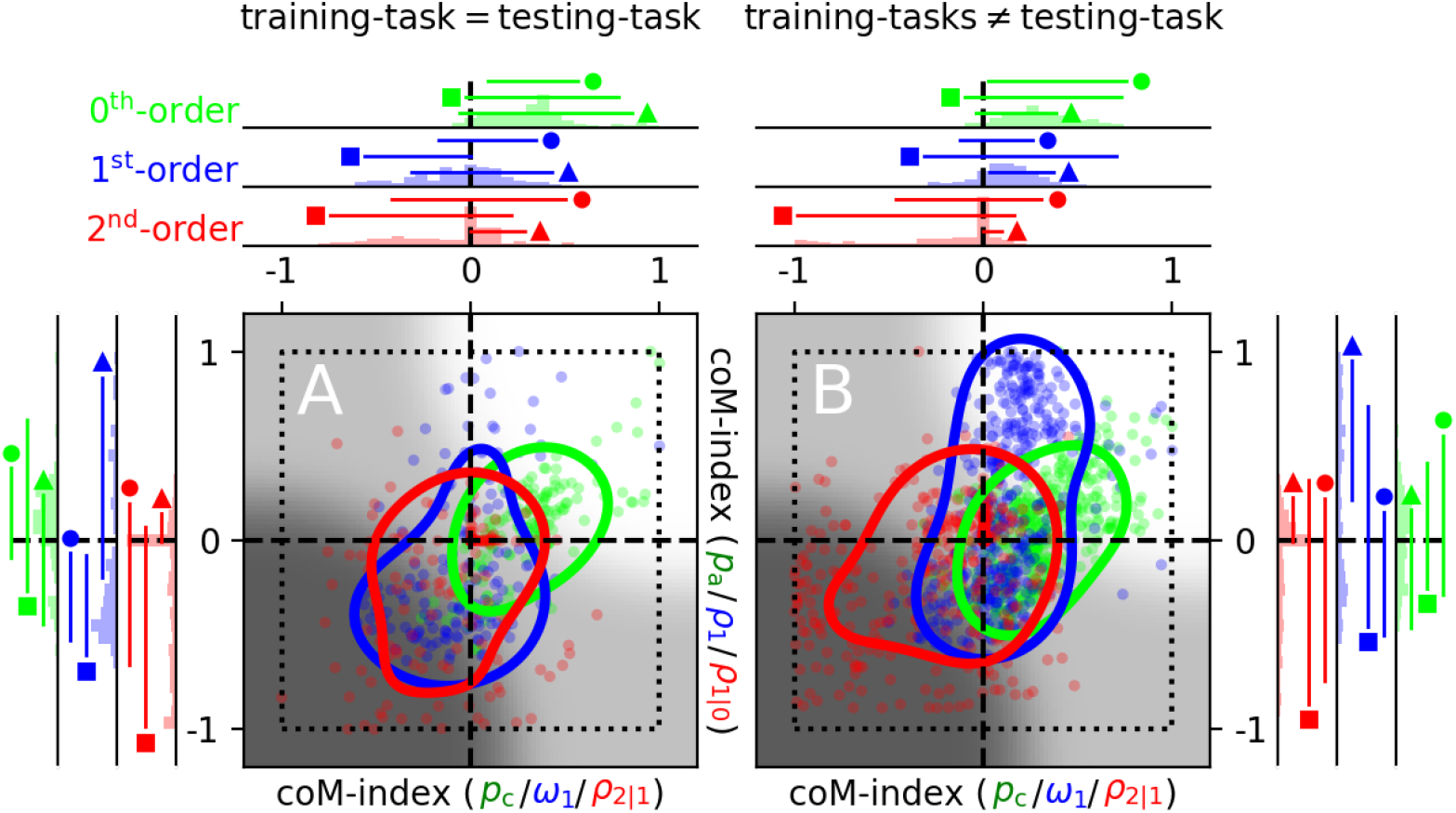
Role of task variety at training. In **Figure 5A**, a given task is adopted for both training and testing phases. In **A** here, two tasks are selected for the training phase, and the remaining task is tested. For example, co-modulation indices for the detection task refer to a network that is trained on polarity-discrimination and energy-extraction tasks. In **B**, all three tasks are combined during the training phase.

## Appendix B

Role of multi-tasking at training

**Figure** .**13A** shows Lenet-5 data across all three tasks, when the network is only trained on one task at a time (e.g. detection) and is then tested on that same task. It collates data from **Figure 5A** (detection), **Figure .12A** (polarity discrimination) and **Figure** .**12C** (energy extraction). **Figure .12A,C** show equivalent data for the other two tasks (polarity discrimination and energy-extraction). **Figure .13B** shows equivalent data when Lenet-5 is trained on two tasks (e.g. detection and polarity discrimination) and tested on the remaining task (e.g. energy extraction), i.e. the network is *not* trained on the task it is subsequently tested on. The results are not substantially different from those obtained by training/testing with the same task.

**Figure 5C** shows the co-modulation plot associated with training Lenet-5 on all three tasks simultaneously. When compared with the co-modulation plots in **Figure** .**13**, this training diet demonstrates slight improvements. In a variant of this protocol, we modified the stimulus to include an explicit task-label by using three different colours for the vertical markers placed above and below each pattern (red lines in **Figure 2**): red for detection, green for polarity discrimination, and blue for energy extraction. This variant produces similar results.

**Figure .14.**
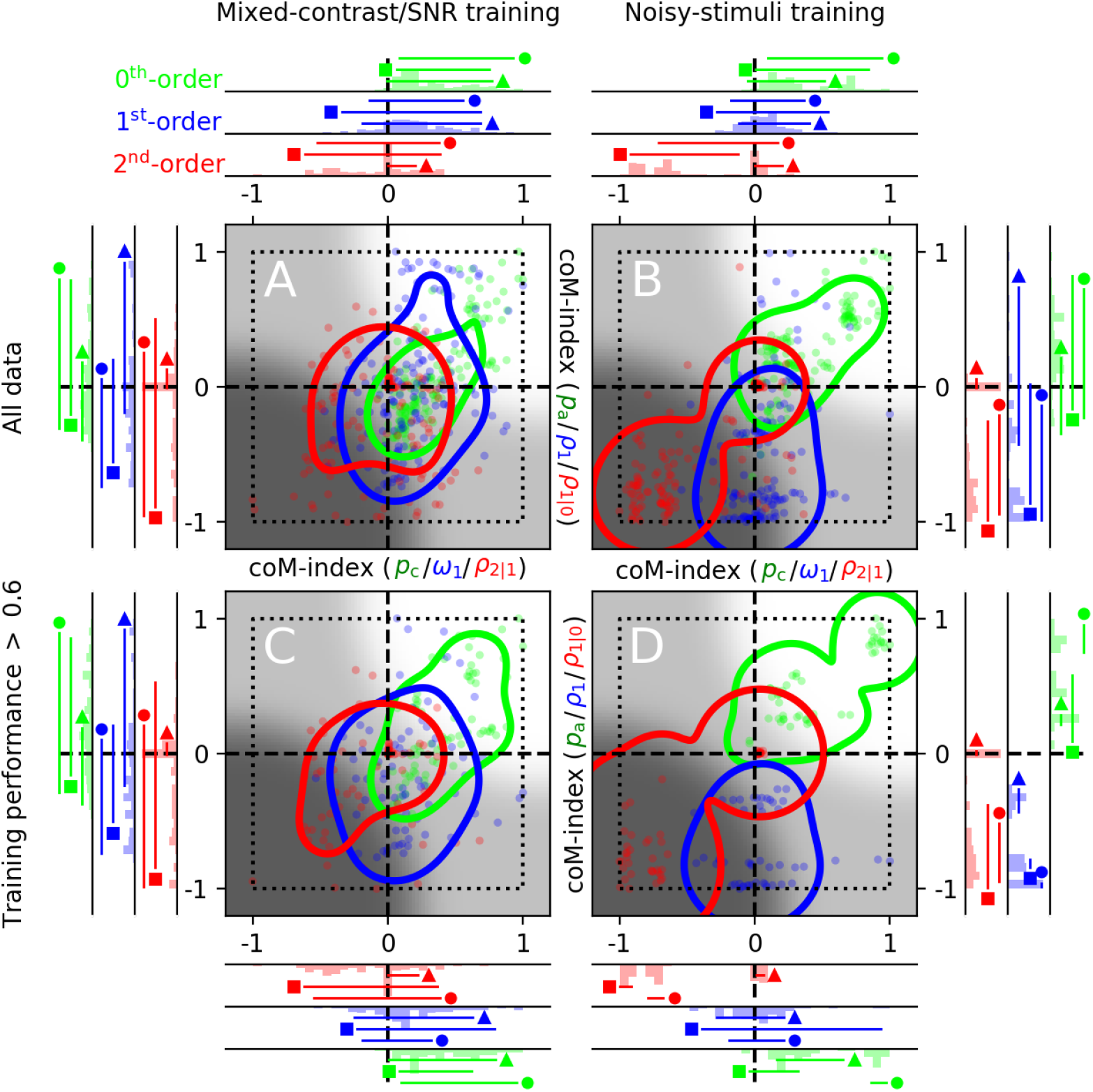
Training with noisy stimuli of varying contrast/SNR. Plotted to the conventions of **Figure .12. A**/**B** refer to Lenet-5 trained on noiseless/noisy stimuli of varying contrast/SNR (training dataset for **B** is statistically identical to testing dataset). **C-D** exclude iterations for which training performance averaged over the last 10 minibatches was below 60% correct.

## Appendix C

Role of stimulus characteristics at training

Section 3.2 refers to simulations for which networks are trained on noiseless high-intensity signals (stimuli to the left of the vertical dashed line in **Figure 2**). This training diet allows networks to quickly learn task-relevant information, while at the same time developing robustness to noise. However, it is conceivable that at least some of the idiosyncratic features displayed by the human data may emerge if networks were trained with noisy stimuli of varying contrast/SNR that are similar to those used during the testing phase.

As a general remark, we observe the following with relation to this possibility: we have performed numerous simulations using noisy training protocols, but we have invariably found that they lead to poorer, not better, results (see also [7]). In larger networks like Resnet-18, good performance is achieved during the training phase but does not transfer to the testing phase (chance performance), indicating that the network is attempting to learn stimulus features that are specific to the noise (and are therefore useless for detecting the target signal). In smaller networks (like Lenet-5), training is degraded by noisy stimuli (more exposure is needed to achieve plateau performance and this remains below 100% correct), with no emergence of first-order/second-order features that convincingly resemble the human data (see below).

As an intermediate step in the transition to the stimulus-set used for testing, we train Lenet-5 with stimuli that are devoid of noise, but vary in contrast/SNR. Technically, stimulus SNR is not defined in the absence of external noise (section 2.1); nevertheless, different SNR levels are associated with different values for the target (central) bar, so stimuli do differ across this dimension despite being noiseless. At the same time, it should be noted that contrast and SNR specifications become confounded in the absence of noise: they both correspond to changes of target bar intensity, without any other concomitant stimulus change. With this in mind, we observe the following. First, plateau performance at training does not always reach perfect performance; for this reason, alongside results from all simulations in **Figure .14A**, we also examine results from simulations for which training performance reaches plateau levels substantially above chance (>60% correct responses) in **Figure .14C**. Although there are some differences between the two co-modulation plots, they confirm the main result demonstrated earlier: network matches human for the zeroth-order characterization, but not for higher-order characterizations.

With the addition of noise at training, the stimulus set adopted to train the network is statistically identical to the stimulus set used to test the network. As shown in **Figure .14B** and **Figure .14D**, this training diet further emphasizes the extent to which different orders of characterization fall within different regions of the co-modulation plot: particularly when the analysis is restricted to simulations associated with better training performance (**Figure.14D**), it is clear that the network is able to reproduce the zeroth-order human trend across all three tasks (green dots fall within the bright region), but it fails at capturing higher orders (blue/red data points).

## References

[1] T. Serre, Deep Learning: The Good, the Bad, and the Ugly, Annu Rev Vis Sci 5 (2019) 399–426.

[2] J. Borowski, C. Funke, K. Stosio, W. Brendel, T. Wallis, M. Bethge, The notorious difficulty of comparing human and machine perception, 2019.

[3] C. Firestone, Performance vs. competence in human-machine comparisons, Proc Natl Acad Sci U S A 117 (43) (2020) 26562–26571.

[4] Z.-Q. Zhao, P. Zheng, S.-T. Xu, X. Wu, Object detection with deep learning: A review, IEEE Transactions on Neural Networks and Learning Systems PP (2019) 1–21. doi:10.1109/TNNLS.2018.2876865.

[5] Y. LeCun, B. Boser, J. S. Denker, D. Henderson, R. E. Howard, W. Hubbard, L. D. Jackel, Backpropagation applied to handwritten zip code recognition, Neural Computation 1 (4) (1989) 541–551.

[6] A. Krizhevsky, I. Sutskever, G. Hinton, Imagenet classification with deep convolutional neural networks, Neural Information Processing Systems 25. doi:10.1145/3065386.

[7] R. Geirhos, D. Janssen, H. Schütt, J. Rauber, M. Bethge, F. Wichmann, Comparing deep neural networks against humans: object recognition when the signal gets weaker.

[8] R. Rajalingham, E. B. Issa, P. Bashivan, K. Kar, K. Schmidt, J. J. DiCarlo, Large-Scale, High-Resolution Comparison of the Core Visual Object Recognition Behavior of Humans, Monkeys, and State-of-the-Art Deep Artificial Neural Networks, J. Neurosci. 38 (33) (2018) 7255– 7269.

[9] R. Geirhos, K. Meding, F. A. Wichmann, Beyond accuracy: quantifying trial-by-trial behaviour of cnns and humans by measuring error consistency (2020). 2006.16736.

[10] A. Doerig, L. Schmittwilken, B. Sayim, M. Manassi, M. H. Herzog, Capsule networks as recurrent models of grouping and segmentation, PLoS Comput Biol 16 (7) (2020) e1008017.

[11] J. Kim, M. Ricci, T. Serre, Not-So-CLEVR: learning same-different relations strains feedforward neural networks, Interface Focus 8 (4) (2018) 20180011.

[12] P. Neri, The empirical characteristics of human pattern vision defy theoretically-driven expectations, PLoS Comput. Biol. 14 (12) (2018) e1006585.

[13] A. A. Faisal, L. P. Selen, D. M. Wolpert, Noise in the nervous system, Nat. Rev. Neurosci. 9 (4) (2008) 292–303.

[14] R. F. Murray, Classification images: A review, J Vis 11 (5) (2011) 1–25 (doi:10.1167/11.5.2).

[15] P. Neri, Stochastic characterization of small-scale algorithms for human sensory processing, Chaos 20 (2010) 045118.

[16] P. Neri, Classification images as descriptive statistics, J. Mathematical Psychology 82 (2018) 26–37.

[17] P. Neri, Visual detection under uncertainty operates via an early static, not late dynamic, non-linearity, Front Comput Neurosci 4 (2010) 151.

[18] J. Kubilius, M. Schrimpf, A. Nayebi, D. Bear, D. L. K. Yamins, J. J. DiCarlo, Cornet: Modeling the neural mechanisms of core object recognition, bioRxivdoi:10.1101/408385.

[19] M. Andreux, T. Angles, G. Exarchakis, R. Leonarduzzi, G. Rochette, L. Thiry, J. Zarka, S. Mallat, J. Andén, E. Belilovsky, J. Bruna, V. Lostanlen, M. J. Hirn, E. Oyallon, S. Zhang, C. Cella, M. Eickenberg, Kymatio: Scattering transforms in python, CoRR abs/1812.11214. 1812.11214. URL http://arxiv.org/abs/1812.11214

[20] M. Ahissar, S. Hochstein, The reverse hierarchy theory of visual perceptual learning, Trends Cogn Sci 8 (10) (2004) 457–464.

[21] F. Wang, J. Huang, Y. Lv, X. Ma, B. Yang, E. Wang, B. Du, W. Li, Y. Song, Predicting perceptual learning from higher-order cortical processing, NeuroImage 124 (2016) 682 – 692.

[22] R. Brunelli, T. Poggio, Template matching: matched spatial filters and beyond, Pattern Recognition 30 (1997) 751–768.

[23] P. Neri, Estimation of nonlinear psychophysical kernels, J Vis 4 (2004) 82–91.

[24] V. Z. Marmarelis, Nonlinear Dynamic Modeling of Physiological Systems, Piscataway: New Jersey: Wiley IEEE Press, 2004.

[25] R. A. Sandler, V. Z. Marmarelis, Understanding spike-triggered covariance using Wiener theory for receptive field identification, J Vis 15 (9) (2015) 16.

[26] D. M. Green, J. A. Swets, Signal Detection Theory and Psychophysics, New York: Wiley, 1966.

[27] D. M. Green, Consistency of auditory detection judgments, Psychol Rev 71 (1964) 392–407.

[28] A. E. Burgess, B. Colborne, Visual signal detection. IV. Observer inconsistency, J Opt Soc Am A 5 (1988) 617–627.

[29] P. Neri, How inherently noisy is human sensory processing?, Psychon Bull Rev 17 (2010) 802–808.

[30] A. J. Ahumada, Classification image weights and internal noise level estimation, J Vis 2 (2002) 121–131.

[31] P. Neri, Nonlinear characterization of a simple process in human vision, J Vis 9 (2009) 1–29.

[32] E. R. Joosten, S. A. Shamma, C. Lorenzi, P. Neri, Dynamic reweighting of auditory modulation filters, PLoS Comput. Biol. 12 (7) (2016) e1005019.

[33] F. Rosenblatt, The perceptron: a probabilistic model for information storage and organization in the brain, Psychol Rev 65 (6) (1958) 386–408.

[34] W. S. Geisler, Ideal observer theory in psychophysics and physiology, Physica Scripta 39 (1989) 153–160.

[35] P. Neri, The elementary operations of human vision are not reducible to template matching, PLoS Comput. Biol. 11 (11) (2015) e1004499.

[36] C. J. McAdams, J. H. Maunsell, Effects of attention on orientation-tuning functions of single neurons in macaque cortical area V4, J. Neurosci. 19 (1999) 431–441.

[37] A. E. Paltoglou, P. Neri, Attentional control of sensory tuning in human visual perception, J Neurophysiol 107 (2012) 1260–1274.

[38] A. Ahumada, R. Marken, Time and frequency analyses of auditory signal detection, J. Acoust. Soc. Am. 57 (1975) 385–390.

[39] C. K. Abbey, M. P. Eckstein, Classification images for detection, contrast discrimination, and identification tasks with a common ideal observer, J Vis 6 (2006) 335–355.

[40] Y. Loewenstein, O. Raviv, M. Ahissar, Dissecting the Roles of Supervised and Unsupervised Learning in Perceptual Discrimination Judgments, J Neurosci 41 (4) (2021) 757–765.

[41] L. K. Wenliang, A. R. Seitz, Deep Neural Networks for Modeling Visual Perceptual Learning, J Neurosci 38 (27) (2018) 6028–6044.

[42] F. Zhuang, Z. Qi, K. Duan, D. Xi, Y. Zhu, H. Zhu, H. Xiong, Q. He, A comprehensive survey on transfer learning, CoRR abs/1911.02685.

[43] L. J. Ba, R. Caruana, Do deep nets really need to be deep?, in: Proceedings of the 27th International Conference on Neural Information Processing Systems - Volume 2, NIPS’14, MIT Press, Cambridge, MA, USA, 2014, p. 2654–2662.

[44] P. Neri, D. M. Levi, Receptive versus perceptive fields from the reverse-correlation viewpoint, Vision Res. 46 (2006) 2465–2474.

[45] W. S. Geisler, Contributions of ideal observer theory to vision research, Vision Res. 51 (7) (2011) 771–781.

[46] J. Cohen, A coefficient of agreement for nominal scales, Educational and Psychological Measurement 20 (1960) 37 – 46.

[47] M. Spilioti, N. Vargesson, P. Neri, Quantitative assessment of intrinsic noise for visually guided behaviour in zebrafish, Vision Res. 127 (2016) 104–114.

[48] D. C. Marr, Vision: A Computational Investigation into the Human Representation and Processing of Visual Information, New York: Freeman, 1982.

[49] M. J. Morgan, Features and the ‘primal sketch’, Vision Res. 51 (2011) 738–753.

[50] K. Kar, J. Kubilius, K. Schmidt, E. B. Issa, J. J. DiCarlo, Evidence that recurrent circuits are critical to the ventral stream’s execution of core object recognition behavior, Nature Neuroscience 22 (2019) 974 – 983.

[51] R. M. Cichy, D. Kaiser, Deep Neural Networks as Scientific Models, Trends Cogn Sci 23 (4) (2019) 305–317.

[52] P. Z. Marmarelis, V. Z. Marmarelis, Analysis of Physiological Systems: the White-Noise Approach, New York: Plenum Press, 1978.

[53] F. E. Theunissen, K. Sen, A. J. Doupe, Spectral-temporal receptive fields of nonlinear auditory neurons obtained using natural sounds, J. Neurosci. 20 (2000) 2315–2331.

[54] A. Fiorentini, L. Mazzantini, Neural inhibition in the human fovea: a study of interactions between two line stimuli, Atti della Fondazione Giorgio Ronchi 21 (6) (1966) 738–747.

[55] C. K. Abbey, M. P. Eckstein, Frequency tuning of perceptual templates changes with noise magnitude, J Opt Soc Am A Opt Image Sci Vis 26 (11) (2009) 72–83.

[56] D. Ringach, R. Shapley, Reverse correlation in neurophysiology, Cognitive Science 28 (2004) 147–166.

[57] P. Neri, Object segmentation controls image reconstruction from natural scenes, PLoS Biol. 15 (8) (2017) e1002611.

[58] Psyphy: A psychophysics driven evaluation framework for visual recognition, IEEE Transactions on Pattern Analysis and Machine Intelligence PP.

[59] A. Doerig, A. Bornet, O. H. Choung, M. H. Herzog, Crowding reveals fundamental differences in local vs. global processing in humans and machines, Vision Res. 167 (2020) 39–45.

[60] T. Xu, O. Garrod, S. H. Scholte, R. Ince, P. G. Schyns, Using psychophysical methods to understand mechanisms of face identification in a deep neural network, in: 2018 IEEE/CVF Conference on Computer Vision and Pattern Recognition Workshops (CVPRW), 2018, pp. 2057–20578.

